# Engineering and characterization of carbohydrate-binding modules to enable real-time imaging of cellulose fibrils biosynthesis in plant protoplasts

**DOI:** 10.1101/2023.01.02.522519

**Authors:** Dharanidaran Jayachandran, Peter Smith, Mohammad Irfan, Junhong Sun, John M. Yarborough, Yannick J. Bomble, Eric Lam, Shishir P.S. Chundawat

**Affiliations:** Department of Chemical and Biochemical Engineering, Rutgers-The State University of New Jersey, Piscataway, NJ 08854, USA; Biosciences Center, National Renewable Energy Laboratory, Golden, Colorado 80401, USA; Department of Plant Biology, Rutgers-The State University of New Jersey, New Brunswick, NJ 08901, USA

**Author notes:** **Corresponding Author:** Shishir P. S. Chundawat.

**Keywords:** Arabidopsis plant protoplasts, Carbohydrate-binding module, Cellulose microfibrils, Cell wall biosynthesis, Confocal laser scanning microscopy, Live-cell imaging, Quartz crystal microbalance with dissipation

## Abstract

- Carbohydrate binding modules (CBMs) are non-catalytic domains associated with cell wall degrading carbohydrate-active enzymes (CAZymes) that are often present in nature tethered to distinct catalytic domains (CD). Fluorescently labeled CBMs have been also used to visualize the presence of specific polysaccharides present in the cell wall of plant cells and tissues.
- Previous studies have provided a qualitative analysis of CBM-polysaccharide interactions, with limited characterization of optimal CBM designs for recognizing specific plant cell wall glycans. Furthermore, CBMs also have not been used to study cell wall regeneration in plant protoplasts.
- Here, we examine the dynamic interactions of engineered type-A CBMs (from families 3a and 64) with crystalline cellulose-I and phosphoric acid swollen cellulose (PASC). We generated tandem CBM designs to determine their binding parameters and reversibility towards cellulose-I using equilibrium binding assays. Kinetic parameters - adsorption (*k_on_*) and desorption (*k_off_*) rate constants-for CBMs towards nanocrystalline cellulose were determined using quartz crystal microbalance with dissipation (QCM-D). Our results indicate that tandem CBM3a exhibits a five-fold increased adsorption rate to cellulose compared to single CBM3a, making tandem CBM3a suitable for live-cell imaging applications. We next used engineered CBMs to visualize *Arabidopsis thaliana* protoplasts with regenerated cell walls using wide-field fluorescence and confocal laser scanning microscopy (CLSM).
- In summary, tandem CBMs offer a novel polysaccharide labeling probe for real-time visualization of growing cellulose chains in living Arabidopsis protoplasts.

## Introduction

Plant cell walls are structurally complex, metabolically dynamic, and extremely rich polysaccharide repositories. These plant cell wall polysaccharides mainly comprise cellulose, hemicelluloses, and pectin, forming diverse networks and extensively interacting with each other (Knox, 2008). Cellulose microfibrils form the major component of the plant cell wall and are intertwined with xyloglucans and other pectic polysaccharides. To access the different polysaccharides, plant cell wall hydrolases contain highly specific, non-catalytic carbohydrate-binding modules (CBMs) along with catalytic domains (CD). These CBMs increase the proximity between CDs and polysaccharides, resulting in efficient enzymatic hydrolysis (Talamantes *et al*., 2016). Different CBMs recognize different polysaccharides based on their amino acid sequences and the topology of the binding site. According to the Carbohydrate-Active enZYmes (CAZy) database (https://www.cazy.org;), there are 94 sequence-based families of CBMs, many of which bind to cell wall polymers (Cantarel *et al*., 2009). For instance, type-A CBM recognizes crystalline polysaccharides such as cellulose, chitin, and mannan. Likewise, type-B CBMs target individual glucan chains, and type-C CBMs bind specifically to small sugars (mono- or disaccharides) (Boraston *et al*., 2004). These CBMs are highly specific, and their specificity depends on the target substrate of its accompanying CD. However, some cellulose-binding CBMs have been reported to be components of enzymes that hydrolyze xylans, mannans, and pectins, other than cellulases (Kellett *et al*., 1990; McKie *et al*., 2001).

In nature, non-catalytic CBMs tend to coexist with other non-catalytic CBMs besides CDs, forming tandem repeats of CBMs. Such tandem CBMs increase the cell wall hydrolases’ overall efficiency by enhancing affinity through prolonged contact with the target substrate (Hashimoto, 2006; Guillén *et al*., 2010; Møller *et al*., 2021). For example, a type-A tandem CBM with three CBM10 domains associated with mannanase showed improved binding and spatial flexibility (Møller *et al*., 2021). Similarly, type-B CBM tandems (CBM17 and CBM28) from *Bacillus* sp. 1139 Cel5 and two family 4 CBMs (CBM4–1 and CBM4–2) from *Cellulomonas* sp. exhibited very tight non-crystalline cellulose binding. Taken individually, these three CBMs recognized different regions of non-crystalline cellulose. (Boraston *et al*., 2003; Kognole & Payne, 2018). Aside from hydrolysis of celluloses, only a few of these tandem CBMs have been engineered to visualize multiple polysaccharides associated with plant cell walls. For example, Herve et al. found that the CBM3a·CBM2b-1-2 tandem constructs bound tightly to cell wall of stem sections where cellulose and xylan were cross-linked closely (Hervé *et al*., 2010). No reports indicate the use of tandem CBMs in imaging single plant protoplasts. Living plant protoplasts provide a distinct advantage as a cell-based system that can be readily used for performing genomics, transcriptomics, proteomics, metabolomics, and epigenetic analyses (Xu *et al*., 2022). This versatile system could be used to characterize cell-wall regeneration and the polysaccharides associated with it. Additionally, regenerating plant cell walls in Arabidopsis contain crystalline and amorphous regions at the surface (Ruel *et al*., 2012). It is imperative that the CBMs’ ability to bind to various forms of cellulose is also characterized critically.

Extensive research has been conducted on CBMs attached to glucanases using bulk biochemical assays (Chundawat *et al*., 2021; Nemmaru *et al*., 2021), single-molecule cellulase motility assays (Brady *et al*., 2015), kinetic modeling (Levine *et al*., 2010; Shang *et al*., 2013), and molecular simulations (Beckham *et al*., 2014; Vermaas *et al*., 2019). However, most of these assays have been conducted on single CBMs attached to fluorescent proteins or glucanases. Consequently, the biochemical behavior of most tandem CBMs under the assays mentioned above has not been studied despite the prevalence of tandem CBMs in nature (Boraston *et al*., 2004; McCartney *et al*., 2006; Hervé *et al*., 2010). Ultimately, the potential of tandem CBMs in the visualization and characterization of cell wall polymers remains mostly unexplored.

Here, using tandem modular constructs, we computed different binding parameters for tandem versus single CBMs. We picked two model type-A CBMs (CBM3a and CBM64) and systematically engineered tandem versions with and without green fluorescent protein (GFP). Using conventional pull-down assays, we calculated the number of binding sites, binding constants, and partition coefficient towards cellulose-I and PASC. Using quartz crystal microbalance with dissipation (QCM - D), we also measured the adsorption and desorption rate constants (*k*_on_ and *k*_off_) for CBM binding towards nanocrystalline cellulose-I. Finally, we used the CBMs generated to visualize the regenerated cell walls in *Arabidopsis thaliana* plant protoplasts to fully understand their potential in real-time visualization of plant cell wall growth using confocal laser scanning microscopy (CLSM) and wide-field fluorescence microscopy.

## Materials and Methods

### Bacterial strains, CBM plasmids, and reagents

Avicel cellulose-I was obtained from Sigma Aldrich under the label Avicel PH-101. Phusion Master Mix was obtained from Thermo Fisher Scientific. Dpn1 and T4 DNA polymerase was obtained from New England Biolabs. Nickel-charged magnetic beads and magnetic racks were purchased from GenScript (NJ). Chemically competent cells were procured from various vendors: *E. cloni* 10g cells from Lucigen (Madison, WI) and *E. coli* BL21-CodonPlus-RIPL [λDE3] from Stratagene (Santa Clara, CA). The pEC-GFP-CBM3a vector was kindly provided by Dr. Brian Fox (University of Wisconsin, Madison, USA) and was used as plasmid backbone to engineer all CBM constructs (Whitehead *et al*., 2017; Chundawat *et al*., 2021). The primers were obtained from Integrated DNA Technologies, and DNA sequencing was performed by Azenta Life Sciences (NJ). All other reagents were purchased from VWR, Thermo Fisher Scientific, and Sigma Aldrich unless mentioned otherwise.

### Synthesis and cloning of wild-type CBM genes into pEC-GFP vector

*E. coli* expression vector pEC-GFP-CBM3a was kindly provided by the Fox lab (UW Madison) (Whitehead *et al*., 2017), which was used as the plasmid backbone to insert the CBM64 gene from *Spirochaeta thermophila*. Cloning and insertion of CBM64 are detailed in our previous work (Nemmaru *et al*., 2021). Plasmid maps for pEC-GFP-CBM3a and pEC-GFP-CBM64 are outlined in Figure S1. Tandem CBMs (pEC-GFP-CBM3a-CBM3a, pEC-CBM3a-CBM3a, pEC-GFP-CBM64-CBM64, and pEC-CBM64-CBM64) were prepared from the original plasmids (pEC-GFP-CBM3a and pEC-GFP-CBM64) using Sequence and Ligation-Independent Cloning (SLIC) protocol (Stevenson *et al*., 2013). The schematic representation of all the CBM constructs is outlined in Figure S2. The SLIC primers used in this study are tabulated in Table S1. The sequences for all CBM constructs generated are summarized in Supplementary Text S1. Briefly, the gene fragment containing the linker and CBM was amplified using polymerase chain reaction (PCR) from the original plasmid available at the Chundawat lab. It was inserted after the ‘GFP-CBM’ region in the original ‘pEC-GFP-CBM’ plasmids to create the ‘tandem CBM’ (also referred to as ‘CBM-CBM’) carrying plasmids. The insert and vector PCR products were purified to remove unreacted nucleotides (deoxynucleoside triphosphates or dNTPs), mixed at optimal ratios, and then transformed into chemically competent *E. cloni* 10G cells to get colonies for screening. The tandem plasmids developed were later used to create non-GFP versions by removing the GFP from those plasmids. All the pEC-GFP-CBM, pEC-GFP-CBM-CBM, and pEC-CBM-CBM plasmids were verified using Sanger sequencing, and the sequence-verified plasmids were stored at −80°C for long-term storage. *E.cloni* 10G cells carrying the plasmid of interest were stored as 15% glycerol stocks and maintained at −80°C.

### Production and purification of recombinant His-tagged CBMs

After sequence verification, all CBM constructs were transformed into *E.coli* BL21-CodonPlus-RIPL [λDE3]. Glycerol stocks were prepared for the transformed strains and were stored at −80°C. These glycerol stocks were later used to inoculate 10 mL Luria Bertani (LB) media in culture tubes containing kanamycin at a 50 μg/ml concentration. This overnight starter culture was used to inoculate 300 mL LB media in 1 L shake flasks containing kanamycin at a 50 μg/mL concentration. The flasks were incubated at 37°C and 200 rpm until the growth reached the exponential phase (OD_600_ ~0.6-0.8). Protein expression was then induced using 0.5 mM IPTG at 25°C for 16 hours. Cells were harvested at 7000x g for 15 mins. 3g of cell pellet was resuspended in 15 mL cell lysis buffer (20 mM phosphate buffer, 500 mM NaCl, 20% (v/v) glycerol, pH 7.4). For every 3g of wet cell pellet, 200 μL protease inhibitor cocktail (1 μM E-64 (Sigma Aldrich E3132)) and 15 μL lysozyme (Sigma Aldrich, USA) was added. Cells were lysed on ice using a Misonix™ sonicator 3000 for 5 mins of total process time at 4.5 output level and pulse settings (pulse-on time: 10 secs and pulse-off time: 30 secs) to avoid sample overheating. The cell lysate carrying the protein of interest was separated from the cell debris by centrifuging at 48,400× g for 45 mins at 4°C. The cell lysate was clarified using a 0.22 μm non-sterile syringe filter after centrifugation. Since all the expressed CBMs contained an N-terminal 8X-HIS tag, the proteins were purified using IMAC (immobilized metal affinity chromatography). The clarified cell lysate was incubated with 2 ml pre-equilibrated Ni^2+^ charged magnetic resin purchased from GenScript. Briefly, the resin was equilibrated with five column volumes of buffer A (100 mM MOPS, 500 mM NaCl, 10 mM Imidazole, pH 7.4). The equilibrated resin was incubated with the clarified lysate at 4°C for 120 mins with gentle agitation. The lysate was removed from the magnetic resin using a magnetic rack. The resin was later washed with five-column volumes of buffer A twice, followed by five-column volumes of (buffer A: buffer B = 95:5). Finally, the proteins were eluted out using five-column volumes of buffer B (100 mM MOPS, 500 mM NaCl, 500 mM Imidazole, pH 7.4). The purified proteins were later desalted into 10 mM MES buffer, pH 6.5. Proteins were also characterized for molecular weights and purity using SDS-PAGE (Figure S3). The concentration of CBM proteins was estimated by measuring the absorbance at 280 nm. The molecular weight and extinction coefficient of all the CBM proteins used in this study are summarized in Table S2. Proteins were later aliquoted, flash-frozen under liquid nitrogen, and stored at −80°C for further assays.

### Generation of PASC from Avicel Cellulose-I

Phosphoric acid swollen cellulose (PASC) was prepared according to the protocol detailed by Zhang et al. (Y.-H. Percival Zhang & Lee R. Lynd, 2006). Briefly, 0.6 ml of distilled water was added to 0.2 g of Avicel cellulose-I (PH-101) to form a wet-cellulose slurry. To this slurry, 10 mL of 86.2% ice-cold phosphoric acid was added with vigorous stirring. The cellulose mixture turned transparent, after which 40 ml of ice-cold distilled water was added at a rate of 10 mL per min. The resulting white cloudy precipitate was removed by centrifugation at 5000× g for 20 mins at 4°C. This step was repeated four more times to remove the phosphoric acid. In addition, 0.5 mL of 2M sodium carbonate was added to neutralize the mixture. Finally, the mixture was washed in ice-cold DI water until the pH was between 5 and 7. The regenerated PASC was stored at 4°C for further binding assays.

### GFP-CBM pull-down binding assays with crystalline cellulose-I and PASC

Binding assays were performed as discussed in previous papers from our group (Chundawat *et al*., 2021; Nemmaru *et al*., 2021). All binding assays were performed with at least six replicates in 300 μL 96-well round-bottomed polypropylene plates (USA Scientific). Each of these replicates was a 200 μL reaction mixture comprising appropriate volumes of CBM dilution to reach effective concentrations of 0–600 μg/mL for obtaining the full binding isotherm. In addition to the CBMs, the mixture comprised 2.5 mg of Avicel cellulose-I, an effective BSA concentration of 2.5 mg/mL to prevent non-specific protein binding, and an effective buffer concentration of 10 mM MES (pH 6.5). To account for protein loss due to denaturation or non-specific binding to the microwells, control reactions without cellulose were also included to obtain total protein concentration. The microplate was then sealed with a 96-well plate mat and incubated inside a USA Scientific hybridization oven at 5 rpm for 60 mins at room temperature with end-over-end mixing. Never-shaken control reactions were also prepared, similar to previously shaken control reactions. This control was used to obtain the calibration curve relating GFP fluorescence and known protein concentration. After incubating the plate for an hour, the microplates were centrifuged at 2000 rpm for 2 mins using an Eppendorf^™^ 5810R centrifuge to separate cellulose from soluble supernatant. Finally, 100 μL of soluble supernatant was picked up from each microwell using a multi-channel micropipette and transferred to black opaque microplates for measuring fluorescence at 480 nm excitation, 512 nm emission with 495 nm cut-off using Molecular Devices^™^ UV spectrophotometer. Schematic representation of the biochemical assay workflow is shown in Figure 1a.

**Figure 1.**
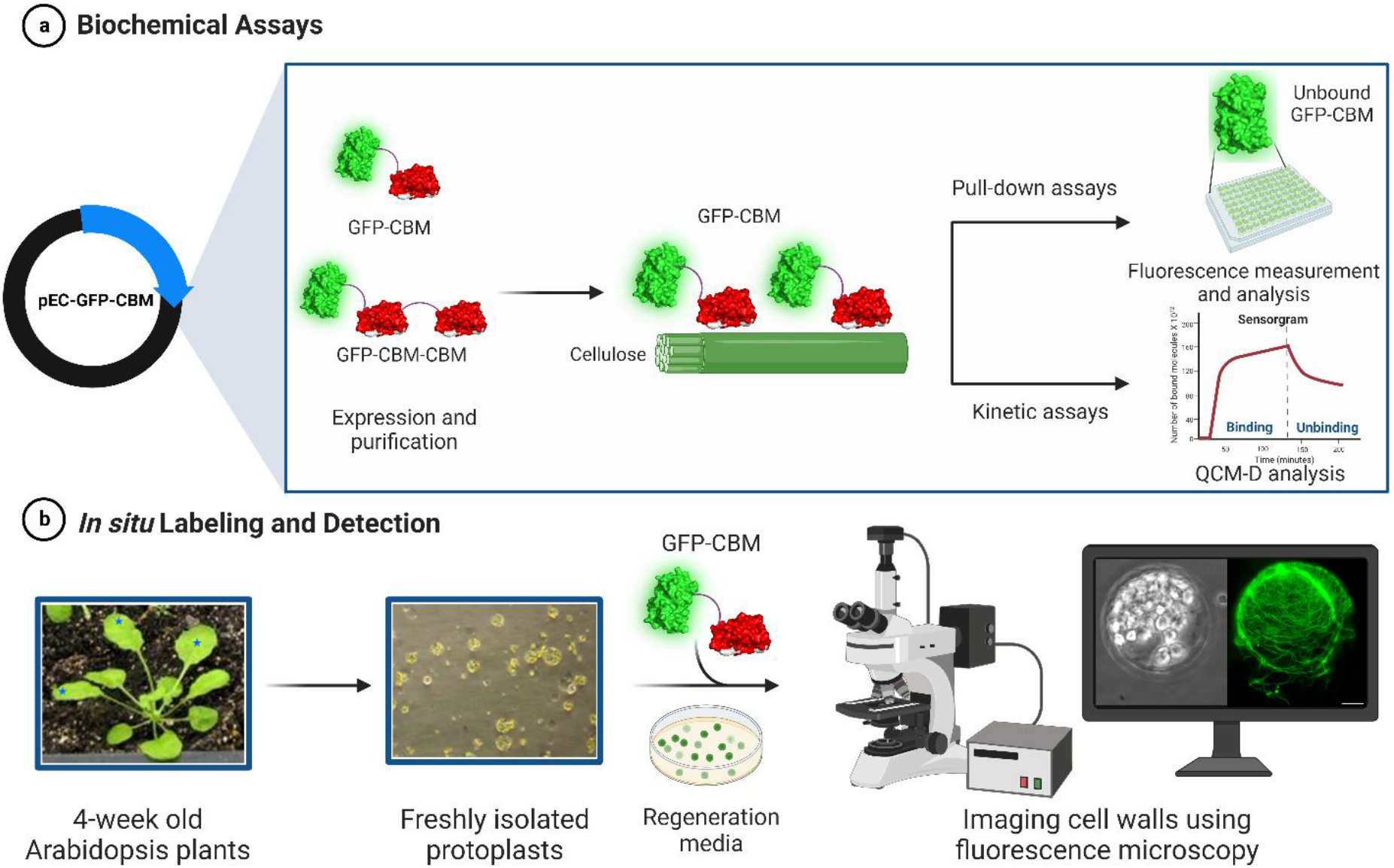
Schematic representation of the workflow in analyzing and utilizing GFP-CBMs to visualize regenerated plant cell walls. (a) pEC-GFP-CBM plasmid used for expressing GFP-CBM3a (*Clostridium thermocellum*) and GFP-CBM3a-CBM3a proteins. Purified GFP-CBMs were used for estimating biochemical and kinetic parameters using pull-down assays and QCM-D analysis. (b) Medium-sized leaves from 4-week-old Arabidopsis plants (marked with blue stars) were used to isolate similarly sized protoplasts. Isolated protoplasts were incubated in the regeneration media along with GFP-CBMs. On the right, regenerated protoplasts with cell walls labeled with GFP-CBM3a as observed under fluorescence microscopy. CBM, carbohydrate-binding module; GFP, green fluorescent protein; QCM-D, Quartz crystal microbalance with dissipation.

### GFP-CBM pull-down binding assay analysis

Preliminary data analysis was performed using Microsoft Excel™ to obtain free protein (μM) and bound protein concentrations (μmol/g cellulose). The data was fit to Langmuir one-site model using the non-linear curve fitting tool in Origin for full-scale binding assays. Curve fitting was done using the Levenberg-Marquardt algorithm with a tolerance of 1e-9.

### Reversibility testing for single and tandem CBM binding to cellulose substrates

A binding reversibility study was performed for all single and tandem CBMs after the completion of the binding assays. After collecting the supernatant for measuring fluorescence in binding assays, 100 μL of reconstitution mixture (2.5 mg/ml BSA + 10 mM MES (pH 6.5)) was added to the remaining original reaction mixtures and shaken controls. The microplate was sealed with a 96-well plate mat and incubated at 5 rpm for 60 mins at room temperature. After incubation, the supernatant was collected, and fluorescence was measured using the spectrophotometer, as mentioned above under the ‘pull-down binding assays’ section.

### Preparation of nanocrystalline cellulose through acid hydrolysis for QCM-D analysis

Nanocrystalline cellulose was prepared from Avicel cellulose-I using the procedure mentioned previously (Nemmaru *et al*., 2021). Briefly, 2 g of Avicel cellulose-I was added to 70 mL 4N hydrochloric acid (HCl) in a glass beaker placed over a water bath maintained at 80°C. The slurry was stirred every 30 mins using a spatula to ensure uniform suspension. After 4 hours, 50 mL of DI water was added to dilute the acid hydrolysis mixture. The slurry was split across 50 mL tubes and centrifuged at 1600x g for 10 mins. The supernatant was discarded, and the pellet was resuspended in 10 mL DI water. This wash step was repeated multiple times until the solution turned hazy and the pH rose to around pH 3.3. The haziness of the supernatant indicated the development of cellulose nanocrystals, and these supernatants were collected for future usage.

### Preparation of cellulose thin films for QCM-D

QCM-D sensors (4.95 MHz quartz crystals, Biolin Scientific QSX-301) were prepared in a manner as described previously (Nemmaru *et al*., 2021). Briefly, quartz crystals were washed thoroughly with deionized water, rinsed with ethanol, and dried with nitrogen gas. The sensors were then placed in a rack and submerged in a solution of 0.02% poly(diallyl dimethyl ammonium chloride), from Sigma Aldrich, for at least 60 mins with orbital mixing (150 rpm) at room temperature. The sensors were then blown-dry once again, followed by spin-coating with 225 μL of cellulose nanocrystal slurry using a Chemat Technology KW-4A spin coater with a pre-cycle spin for 3 secs at 1500 rpm, followed by a spin cycle for 60 secs at 3000 rpm. This spin coating step is repeated 4-8 times to obtain a uniform cellulose film thickness of ~20-40 nm (as measured using the QSoft software using the Sauer brey model and an assumed density of 1191 kg/m^3^). Note that the number of necessary cellulose spin coating steps is dependent on the concentration of cellulose nanocrystals in the prepared slurry and may need to be optimized on a case-to-case basis.

### Quartz crystal microbalance with dissipation (QCM-D) based CBM-cellulose binding assay

Binding and unbinding assays were performed on a QSense E4 instrument (Nanoscience Instruments). The quartz sensors with a deposited cellulose film were loaded into the instrument and allowed to equilibrate overnight in 10 mM MES buffer pH 6.5. Following the overnight incubation, the frequency and resonance changes associated with harmonics 1, 3, 5, 7, and 9 were tracked and monitored for stability for at least 5 mins in the MES buffer prior to loading CBMs. All CBM proteins were diluted to a concentration of 5 μM and passed over the sensors at a flow rate of 100 μL/min for at least 10 mins or until saturation was observed. Unbinding of proteins was tracked by flowing 10 mM MES buffer pH 6.5 at 100 μL/min for at least 20 mins. The sensors and Qsense chambers were then rinsed with 5% Contrad solution followed by deionized water to remove any traces of residual protein. The frequency and dissipation traces were analyzed using an in-house data analysis routine based on binding and unbinding equations derived previously (Nemmaru *et al*., 2021).

### Analysis of QCM-D-based binding assay data to obtain kinetic parameters

Binding curves of QCM-D data were analyzed using RStudio based on equations derived previously (Nemmaru *et al*., 2021). Data corresponding to the frequency changes of the third harmonic of each experiment was transformed with the Sauerbrey equation to produce a change in mass (ng.cm^-1^) associated with CBM binding and un-binding. The area of the sensor gaining mass was found to have a radius of 0.5 cm, and this value was used to determine the total ng of protein binding/unbinding to the cellulose surface.

It was noted that not all the mass that is added to the cellulose sensor is removed during the unbinding experiments, and that some CBM appears to remain bound even after extensive washing. We postulate that this may be due either to a difference in the nature and interaction of CBMs with the nanocellulose coating the surface compared to Avicel-based cellulosic materials in other pull-down experiments, or perhaps due to incomplete coating of the sensor surface with nanocellulose, allowing some CBM to bind irreversibly to the underlying poly(diallyl dimethylammonium chloride) layer. To accommodate this finding and provide a more suitable fit for the unbinding equation, the equation used to fit the unbinding data included a term (CBM_remain_) which accounted for the irreversible CBM binding and represented the number of CBM molecules still bound to the sensor surface after washing. The inclusion of this term produced the following equation, which was used to fit the data.

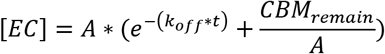

[EC] - Number of binding sites occupied by protein (μmol)
A - Number of available binding sites on nanocellulose film
*k_off_* - Dissociation rate constant (min^-1^)
t - time (min)
CBM_remain_ - Number of CBM molecules that remain bound after washing

### Preparation of plant samples for imaging plant protoplasts

*Arabidopsis thaliana* Col-0 (Columbia) was used as plant material for all experiments. *Arabidopsis thaliana* mesophyll protoplasts were isolated as described in detail by Yoo et al (Yoo *et al*., 2007). Briefly, 3–4-week-old, uniformly sized leaves were selected to obtain consistently sized protoplasts (Figure 1b). Thin leaf strips of 0.5-1 mm thickness were cut off from the middle portion of the leaves. These leaf strips were immediately transferred to an enzyme solution (0.4 M Mannitol, 20 mM KCl, 20 mM MES, 1.5% Cellulase R10 (Yakult, Japan), 0.4% Macerozyme R10 (Yakult, Japan), 10 mM CaCl_2_, 5 mM 2-mercaptoethanol, 0.1% BSA). The leaf strips were vacuum infiltrated to perfuse it with the enzyme solution and then incubated in the dark for 3-5 hours to digest the cell wall completely. The protoplasts were then diluted with an equal volume of sterile W5 solution (154 mM NaCl, 125 mM CaCl_2_, 5 mM KCl, and 2 mM MES in ultrapure water). The undigested leaf material was filtered out using a pre-washed 75 μm nylon mesh. Protoplasts were then collected by centrifugation at 1000 rpm for 3 mins at 4°C. The residual enzymatic solution was removed by washing the protoplast cells with 10 mL of W5 solution twice. After washing, the protoplasts were resuspended in 0.5 mL of W5 solution. The quality of the protoplasts was observed under a simple light microscope (Figure 1b). The concentration of protoplasts was measured using a hemocytometer and then adjusted to 2 × 10^5^ protoplasts mL^-1^ of W5 solution. 200 μL of protoplasts at 2 × 10^5^ concentration was incubated in cell wall regeneration media containing an equal volume of WI solution (0.5 M Mannitol, 4 mM MES, 20 mM KCl) and 2M2 media (Gamborg’s B-5 basal medium with minimal organics 6.4g/L (Sigma), 0.8M Trehalose, 0.1 M Glucose, 2 μM 3-naphthalene acetic acid (NAA) pH 5.7). The protoplasts were incubated for 17 hours at room temperature under a Philips hue lamp. After incubation, the regenerated protoplasts were isolated by spinning at 300x g for 3 mins at 4°C. The isolated protoplasts were then fixed by incubation on ice for 10 mins in 200 μL of 1% glutaraldehyde. The protoplasts were then washed with 250 μL of 12% sorbitol twice. Samples were resuspended in 50 μL of 12% sorbitol and stored on ice until use.

### Fluorescence and Confocal Laser Scanning Microscopy

Calcofluor white solution (100 μL of 0.001%) was added to 200 μL of regenerated protoplasts and incubated at room temperature for 5 mins. Excess calcofluor solution was removed after spinning down the cells at 1000 rpm for 2 mins. The stained cells were washed with 12% sorbitol solution twice. 10 μL of the regenerated protoplast samples were placed on a glass slide that was layered with a coverslip. Calcofluor-stained cell fibers were visualized under the DAPI channel in Olympus FSX 100 microscope. Staining using CBMs was done in a similar manner, where the final concentration of CBMs in the solution was 100 nM. Excess CBMs were removed, and the cells were observed under the GFP channel for GFP-CBMs and the red channel for DSR-CBMs.

For CLSM, the samples were prepared as mentioned above, and the glass slides were then placed onto the inverted platform of a Zeiss LSM 710 confocal microscope, and the cells were imaged under the 488-nm channel for GFP CBMs and 561 nm channel for DSR-CBMs. All images were processed with Zeiss imaging software, ImageJ, or Adobe Photoshop.

## Results

### Type-A tandem CBMs show reduced binding affinity compared to single CBM counterparts

We prepared two type-A tandem CBMs, CBM3a (*Clostridium thermocellum*) (Lehtiö *et al*., 2003) and CBM64 (*Spirochaeta thermophila*) (Schiefner *et al*., 2016; Pires *et al*., 2017) by fusing the linker-CBM region to the N-terminal region of GFP-CBM as shown in Figure S2. Additionally, we studied the binding and activity of GFP-CBM and GFP-CBM-CBM tandem constructs toward Avicel cellulose-I and PASC, respectively (Figure 2a; Figure S4-5).

**Figure 2.**
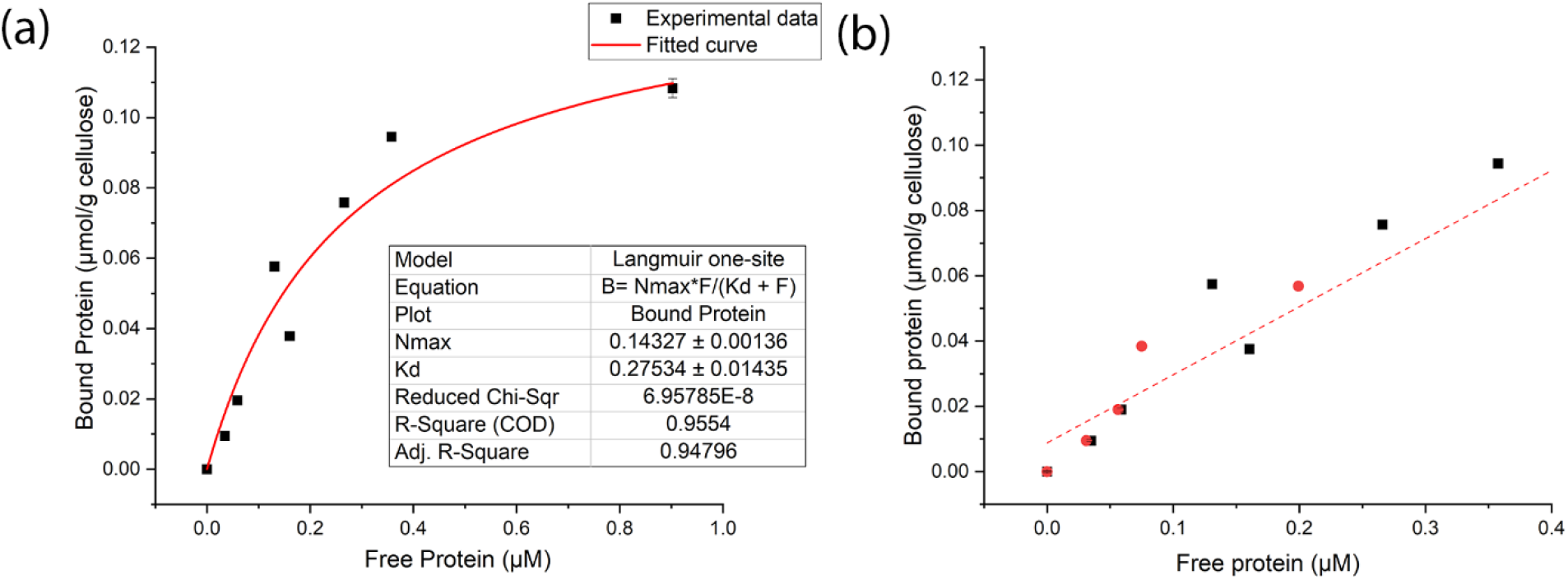
(a) Binding curve of GFP-CBM3a towards cellulose-I using Langmuir-type adsorption model, (b) Binding reversibility of GFP-CBM3a binding towards cellulose-I; Black and red dots represent original and re-equilibrated data points; Red line represents linear fit joining original data points.

First, we performed fluorescence-based pull-down binding assays for GFP-CBM3a and GFP-CBM64 – single and tandem versions against Avicel cellulose-I at varying protein concentrations (ranging between 0 and 600 μg/ml). The resulting data were fit to Langmuir isotherm one-site models to obtain the maximum number of binding sites (N_max_) on the substrate, binding dissociation constant (K_d_), and partition coefficient (see Table 1). The number of available binding sites was the maximum for GFP-CBM3a compared to other CBM versions. Regardless of the substrate (cellulose-I or PASC), GFP-CBM3a always showed a significant similarity in binding affinity (i.e., the inverse of K_d_). The binding affinity was also observed to be higher for GFP-CBM3a compared to GFP-CBM3a-CBM3a against cellulose-I. Nevertheless, the binding affinity parameters are comparable between the single and tandem versions of CBM64. The order of affinity towards cellulose-I is GFP-CBM3a~GFP-CBM64~GFP-CBM64-CBM64>GFP-CBM3a-CBM3a>GFP-CBM17.

**Table 1.** Binding parameters N_max_, K_d_ for GFP-CBM3a, GFP-CBM3a-CBM3a, GFP-CBM64, GFP-CBM64-CBM64, and GFP-CBM17 obtained from Langmuir one-site model fitted to full-scale binding assay data. Errors are standard deviations of the mean obtained from the fitting analysis. Each experiment was performed using six replicates for each protein concentration.

| Construct | Cellulose-I |  |  | PASC |  |  |
| --- | --- | --- | --- | --- | --- | --- |
| | $N_{\max}$<br>($\mu\text{mol/g}$ cellulose) | $K_d$ ($\mu\text{M}$) | $N_{\max}/K_d$ (L/g cellulose) | $N_{\max}$<br>($\mu\text{mol/g}$ cellulose) | $K_d$ ($\mu\text{M}$) | $N_{\max}/K_d$ (L/g cellulose) |
| GFP-CBM3a | 0.14±0.00 | 0.28±0.01 | 0.51±0.09 | 0.17±0.00 | 0.30±0.02 | 0.57±0.04 |
| GFP-CBM3a-CBM3a | 0.09±0.00 | 1.29±0.26 | 0.07±0.02 | 0.06±0.00 | 0.29±0.00 | 0.21±0.10 |
| GFP-CBM64 | 0.02±0.00 | 0.29±0.03 | 0.07±0.03 | 0.13±0.01 | 2.12±0.53 | 0.06±0.02 |
| GFP-CBM64-CBM64 | 0.04±0.00 | 0.30±0.03 | 0.13±0.10 | 0.11±0.00 | 0.48±0.04 | 0.23±0.06 |
| GFP-CBM17 | 0.09±0.00 | 5.20±0.35 | 1.71±0.18 | 0.10±0.02 | 0.93±0.27 | 0.11±0.07 |

Similarly, we subjected all the CBM constructs to binding assays against PASC. We observed a 7-fold and a 1.25-fold increase in the number of available binding sites (N_max_) for CBM3a compared to CBM64 towards cellulose-I and PASC, respectively. Similar to cellulose-I, there was no significant difference in the number of binding sites (N_max_) available for GFP-CBM64, GFP-CBM64-CBM64, and GFP-CBM17. Nevertheless, there was a ~3-fold reduction in N_max_ for the tandem CBM3a compared to the single CBM3a. However, the binding affinity trend towards PASC was found to be similar for single and tandem CBM3a. GFP-CBM17 was found to have the maximum binding affinity towards PASC among all other CBMs, consistent with previous studies (McLean *et al*., 2002).

Furthermore, the partition coefficients of single CBM3a against cellulose-I and PASC were nearly 7- and 2-fold higher than the tandem CBM3a. This reduction in partition coefficient for tandem CBMs on PASC is probably due to the significant steric clashes it encounters with the non-native surface of amorphous cellulose. The uneven topology of the hydrophobic binding face probably impairs the accessibility for tandem CBMs (Chundawat *et al*., 2021; Nemmaru *et al*., 2021). On the other hand, the GFP-CBM64 has a 2- and a 4-fold reduction in partition coefficient compared to tandem CBM64. GFP-CBM17 is highly specific to amorphous cellulose and showed a significant reduction (~5-fold) in binding affinity and an increased partition coefficient towards cellulose-I.

The reversibility of tandem CBMs has not been studied in detail in the past, and most of the reported binding reversibility data for single CBMs has been contradictory (Lim *et al*., 2014; Møller *et al*., 2021). Here, we showed that the binding of single and tandem GFP-CBM3a is reversible on both cellulose-I and PASC ruling out the possibility of protein structural deformation on the cellulose surface (Figure 2b; Figure S6). However, GFP-CBM64 is irreversible on both cellulose-I and PASC, similar to our previous studies (Nemmaru *et al*., 2021). Tandem CBM64 is irreversible only towards PASC and not towards cellulose-I (Figure S7). Conversely, GFP-CBM17 is reversible on both PASC and cellulose-I, respectively (Boraston *et al*., 2003; Blake *et al*., 2006).

### QCM-D assays show a reduced binding behavior for tandem CBMs

Methods describing cellulose film preparation, QCM-D binding assays, and data analysis are discussed in detail in the ‘Materials and Methods’ section. To measure the binding kinetics of single and tandem CBMs on nanocrystalline cellulose, QCM-D binding assays were prepared (Figure 3a). Briefly, the number of protein molecules bound to the cellulose film was calculated by converting the frequency data obtained from the binding and unbinding of proteins (Brunecky *et al*., 2020). Sauerbrey equation was used to obtain the mass of adsorbed protein on cellulose film using the frequency change (Kankare, 2002). In particular, the unbinding regime was fitted to an exponential decay to obtain the true desorption rate (*k_off_*) as shown in Figure 3b. The effective adsorption rate constant (*nk_on_*) was measured using desorption rate *k_off_* from the QCM-D analysis and *N_max_, K_d_* obtained from the equilibrium binding assay results. These kinetic parameters were chosen since they were found to be an integral part of the kinetic models that we had reported previously (Gao *et al*., 2013; Nemmaru *et al*., 2021). QCM-D analysis shows a ~1.7-fold increase in *k_off_* for GFP-CBM3a-CBM3a compared to GFP-CBM3a, indicating a reduced binding affinity for tandem CBM3a compared to the single one. However, the *nk_on_* value for tandem CBM3a increased 5-fold compared to the single CBM3a. Conversely, a similar trend was observed for the *k_off_* values between tandem and single versions of CBM64 towards cellulose-I nanocrystals (Figure 3c). To account for any effects in *k_off_* and *nk_on_* values arising from GFP being fused with CBM3a, we also prepared tandem CBM3a and single CBM3a without GFP. Interestingly, the non-GFP version of CBM3a-CBM3a showed a reduction in the *k_off_* value (nearly 5-fold) compared to the GFP version (Figure S8; Table 2), suggesting that GFP fusion may increase the off-rate for the tandem CBM3a dimer. On the other hand, CBM3a seemed to have an increased off-rate when GFP was removed.

**Figure 3.**
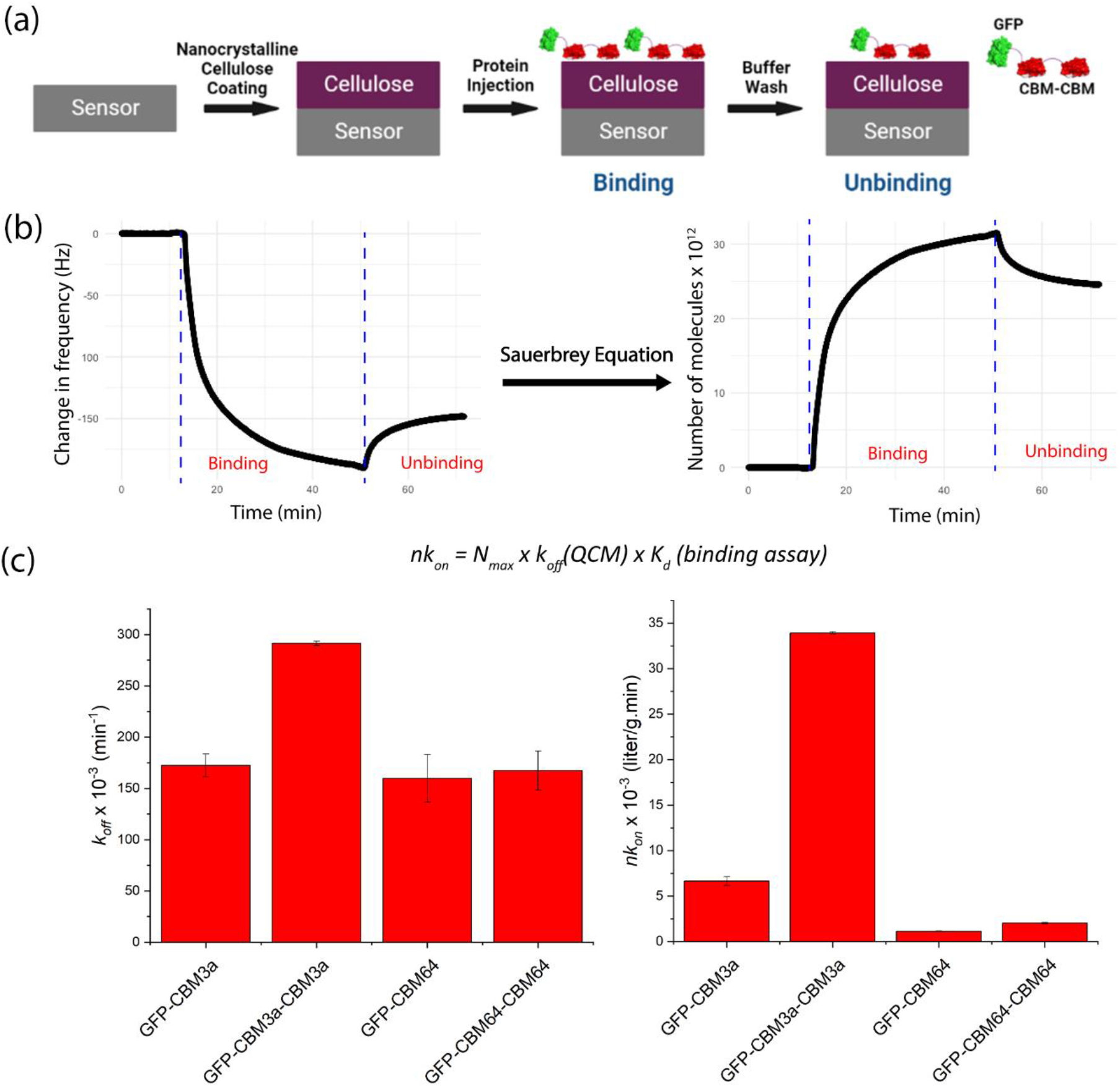
(a) Schematic representation for QCM-D-based tandem CBM - nanocrystalline cellulose-based binding assay. (b) Frequency (Hz) versus time (min) data for a representative protein (GFP-CBM3a) was converted to a sensorgram (number of protein molecules × 10^12^ versus time (mins)) using the Sauerbrey equation. The binding and unbinding data in the plot on the right were then fitted to an exponential rise and decay function, respectively, as described in detail under the methods section. (c) *nK_on_* (right) was calculated using the formula stated, and *nK_on_*, and *K_off_ (left)* for GFP-CBM3a, GFP-CBM3a-CBM3a, GFP-CBM64, and GFP-CBM64-CBM64 toward cellulose-I.

In summary, GFP-CBM3a showed a reduced *nk_on_* compared to GFP-CBM3a-CBM3a. Removing GFP reduced the off-rate drastically for tandem CBM3a but increased it for single CBM3a. On the other hand, CBM64 didn’t have a significant difference in their *k_off_* values. These results suggest that there is no significant difference in binding affinity for single and tandem versions of CBM64 towards cellulose-I. Also, a weaker binding is probably more prevalent in the case of GFP tandem CBM3a towards cellulose-I, which could be determined from a higher desorption rate which is overcome by removing GFP. The *k_off_, nk_on_*, and number of bound molecules are summarized in table 2.

**Table 2.** Kinetic rate constants for GFP-CBM3a, GFP-CBM3a-CBM3a, GFP-CBM64, GFP-CBM64-CBM64, CBM3a, and CBM-CBM3a adsorption and desorption toward nanocrystalline cellulose allomorphs estimated using QCM-D-based binding assay data. n.a. – Not available.

| | $k_{off} \times 10^{-3} (min^{-1})$ | $nk_{on} \times 10^{-3} (L.g^{-1}.min^{-1})$ | $A (x10^{-12} \text{ bound molecules})$ |
| --- | --- | --- | --- |
| GFP-CBM3a | 172.45±11.11 | 6.65±0.47 | 27.91±2.26 |
| GFP-CBM3a-CBM3a | 291.56±2.02 | 33.92±0.00 | 18.20±2.55 |
| GFP-CBM64 | 159.86±23.24 | 1.12±0.00 | 17.89±1.70 |
| GFP-CBM64-CBM64 | 167.35±19.02 | 2.03±0.00 | 16.32±0.00 |
| CBM3a | 223.12±1.86 | n.a. | 23.90±0.42 |
| CBM3a-CBM3a | 64.24±16.84 | n.a. | 30.15±0.70 |

QCM-D has been utilized as a tool to study the binding of full-length cellulases to different types of cellulose (Brunecky *et al*., 2020), lignin, and pretreated biomass (Kumagai *et al*., 2014; Haarmeyer *et al*., 2017). QCM-D has also been used to study CBMs associated with oxidases (Mollerup *et al*., 2016) and cellulase-hemicellulase complexes (Freelove *et al*., 2001a). These reports were mostly confined to single wild-type and engineered CBMs associated with cellulases and other hydrolytic enzymes. We have previously examined the viscoelastic properties of CBM-cellulose binding using QCM-D analysis (Chundawat *et al*., 2021; Nemmaru *et al*., 2021). Here, we explored the kinetic constants of tandem CBMs using QCM-D to identify if they have an improved binding affinity and if they could be used as a weak or tight-binding imaging probe in plant cells. A similar trend in adsorption and desorption rate constants was observed in CBM64 for both tandem and single versions suggesting no apparent avidity advantage for tandem CBM64. Similar behavior was observed in SusF, a starch-utilizing system with tandem CBMs which didn’t show an improved affinity compared to its single and mutated versions (Cameron *et al*., 2012). In contrast, GFP tandem CBM3a showed a higher desorption rate and, ultimately, a lower affinity towards Avicel cellulose-I. This reduced affinity towards crystalline cellulose could be attributed to the linker length that connects the GFP to the CBM3a as well as between the two CBMs present in tandem CBM constructs. The native linker used in this study is 42 amino acids long and has been found to be highly flexible in our previous study (Bandi *et al*., 2020). Compared to shorter linker lengths, the inclusion of a flexible linker resulted in a reduced activity when fused to a cellulase (CelE from *Clostridium thermocellum)*. Similar behavior was observed when double CBMs were fused to cellulases, Cel6A and Cel7A (from *Trichoderma reesei*). The 48 aa-linker decreased the binding affinity and capacity of the tandem CBM compared to reduced linker lengths (Arola & Linder, 2016).

### Fluorescence microscopy for detecting cellulose fibrils during protoplast cell wall regeneration

Procedures for preparing Arabidopsis wild-type mesophyll protoplasts and cell wall regeneration conditions have been described in detail under the ‘Materials and Methods’ section. The isolated protoplasts were healthy and relatively homogeneous in size, as observed under a simple light microscope (Figure 1). After incubation of protoplasts in regeneration media (WI+2M2), the cells were fixed and stained with calcofluor, a β-glucan-specific dye. Calcofluor white has been traditionally used to obtain high-resolution fluorescent images of cellulose without autofluorescence (Kuki *et al*., 2017). After 17 h of incubation in the regeneration media, the cell wall network was spread over the entire surface of the plant protoplast (Figure S9a). However, calcofluor is also known to bind to callose - a β-1,3 glucan present on the plant surface. Besides non-specific binding targets, calcofluor is also known to affect the *in vivo* assembly of cellulose microfibrils (Haigler *et al*., 1980). Hence, the addition of calcofluor to the cell wall regeneration media could potentially induce cytotoxicity for plant cells and preclude its use in monitoring cell wall regeneration in live protoplasts.

To overcome the cytotoxicity, improve the specificity, and predominantly visualize cellulose network around regenerated plant protoplasts, engineered CBMs could be ideal probes to be employed for imaging. 100 nM GFP-CBM3a was used to visualize the regenerated cell wall under fluorescence microscopy. Both calcofluor and GFP-CBM3a stained cells showed the presence of fibrous structures on the surface of protoplasts after overnight incubation in regeneration media. The developed fibers labeled by GFP-CBM3a were distinct and clear when observed under the green emission channel using a fluorescence microscope (Figure S9b).

### Type-A CBMs facilitate Arabidopsis thaliana cell wall visualization

To explore further the general significance of the data mentioned above, the other type-A CBMs analyzed biochemically were also used to image regenerated cell walls of plant protoplasts. All the CBMs used bound to the cell wall surface of protoplasts, typical of type-A CBMs. Confocal laser scanning microscopy (CLSM) was used to acquire images with high-resolution and across multiple focal planes. CLSM images showed that the regenerated plant protoplasts had a pronounced accumulation of fluorescence along the edges of the cell. Both GFP-CBM3a and GFP-CBM3a-CBM3a bound to fibers on the cell wall compared to the control with no CBMs (Figure 4; Supplementary videos 1-4). To check if other type-A tandem CBMs behave in a similar fashion, GFP-CBM64-CBM64 was used to image the regenerated plant protoplasts. Cell wall fibers were observed throughout the edges of the protoplast (Figure S10; Supplementary videos 5). Alternatively, the DS-red versions of CBM3a also showed similar probing characteristics (Figure S11; Supplementary videos 6-7). However, the autofluorescence arising from plant chloroplasts under the red channel could also contribute to the signals arising from DS-red fluorescence (Krause & Weis, 1991). Hence, GFP-CBMs were used for further experiments.

**Figure 4.**
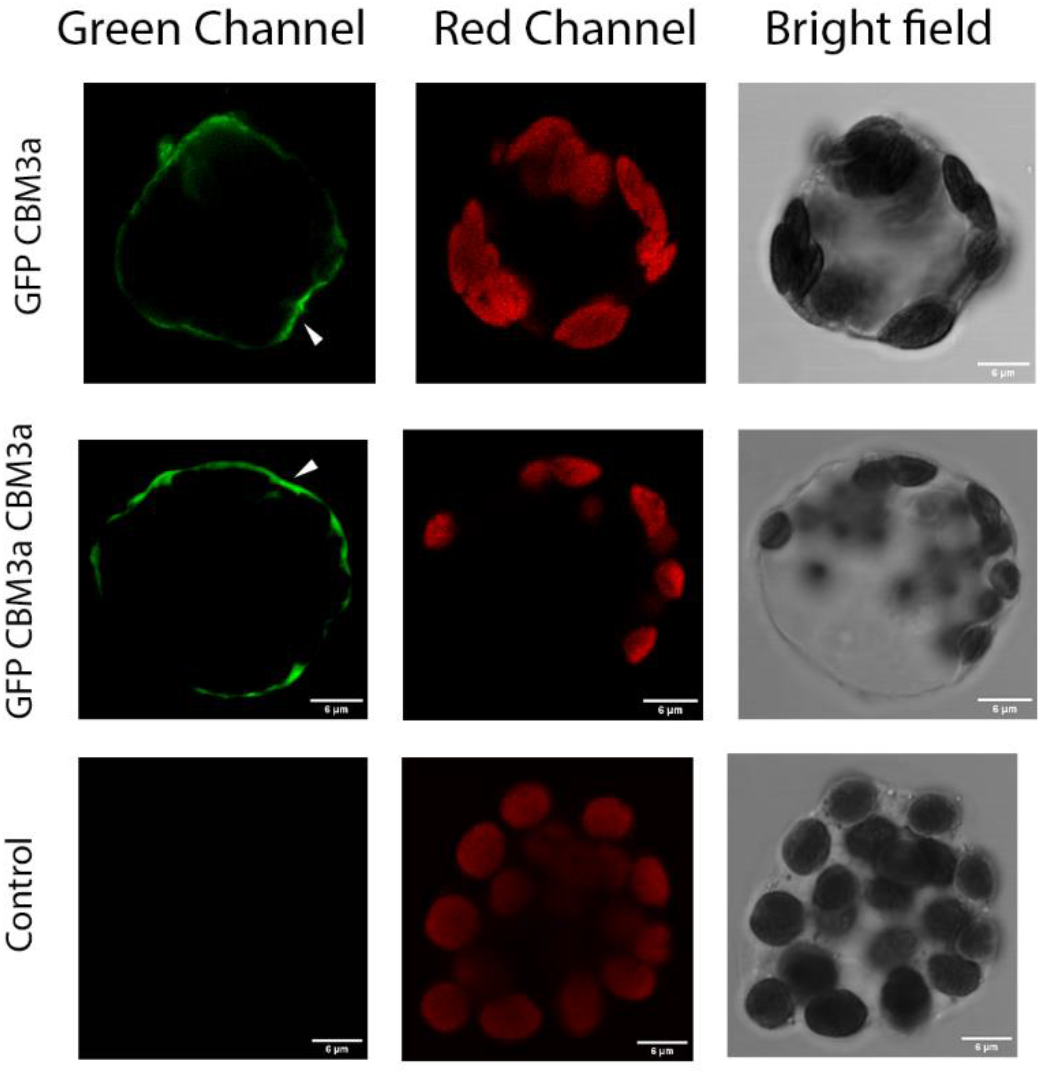
Confocal laser scanning microscopy images of GFP-CBM3a and GFP-CBM3a-CBM3a bound to regenerated plant cell walls of Arabidopsis mesophyll protoplasts. The cell wall labeled with GFP-CBM is visible under the green channel (left), and chloroplasts are visible under the red channel due to autofluorescence (middle) and the bright field image of the intact protoplasts (right). Control had no GFP-CBM added before imaging. White arrows indicate the GFP-CBMs binding to regenerated cell walls. (Scale bars, 6 μm.)

### GFP-CBM3a probe potentiates real-time imaging of cellulose synthesized by plant protoplasts

Previous studies on imaging of plant cell walls were performed using immunocytochemistry and indirect immunofluorescence (McCartney *et al*., 2006; Knox, 2012). In particular, the time course of imaging of plant cell walls was performed by isolating regenerated protoplasts at specific time intervals and labeling them with calcofluor (Kuki *et al*., 2017) or by directly visualizing plant roots using S4B dye (Anderson *et al*., 2010). These studies fail to capture the continuous dynamics of cellulose growth and movement on the surface of regenerating plant protoplasts. These dyes are also toxic and cannot be added to the regeneration media but only to fully developed cells. The absence of molecular probes that could be readily added to the regeneration media is one of the primary reasons for the lack of complete understanding of cellulose regeneration kinetics and other dynamic properties. Conversely, type-A CBMs are specific to cellulose and are non-toxic to plant cells which makes them attractive to overcome this bottleneck.

To examine if these CBMs are compatible with live protoplasts and thus may not interfere with the cell wall regeneration process, we added 100 nM of GFP-CBM3a to the regeneration media (WI+M2) from the outset of the regeneration process. For controls, we had protoplasts in regeneration media with 100 nM GFP and no GFP or GFP-CBM3a. Firstly, as mentioned previously, the regeneration of protoplasts was confirmed by adding GFP-CBM3a to the regenerated protoplasts. Cell wall fibers were clearly distinguishable and visible under fluorescence microscopy (Figure 5a). Similarly, protoplasts incubated in regeneration media, along with GFP-CBM3a, also exhibited a uniform distribution of cellulose at the cell surface compared to cells incubated with only GFP (Figure 5b). Taken together, these results show that GFP-CBMs could be readily added to the regeneration media to continuously monitor the growing cellulose chains. Such a system should now provide the tools to monitor cell wall growth and dynamics in real-time with live cells.

**Figure 5.**
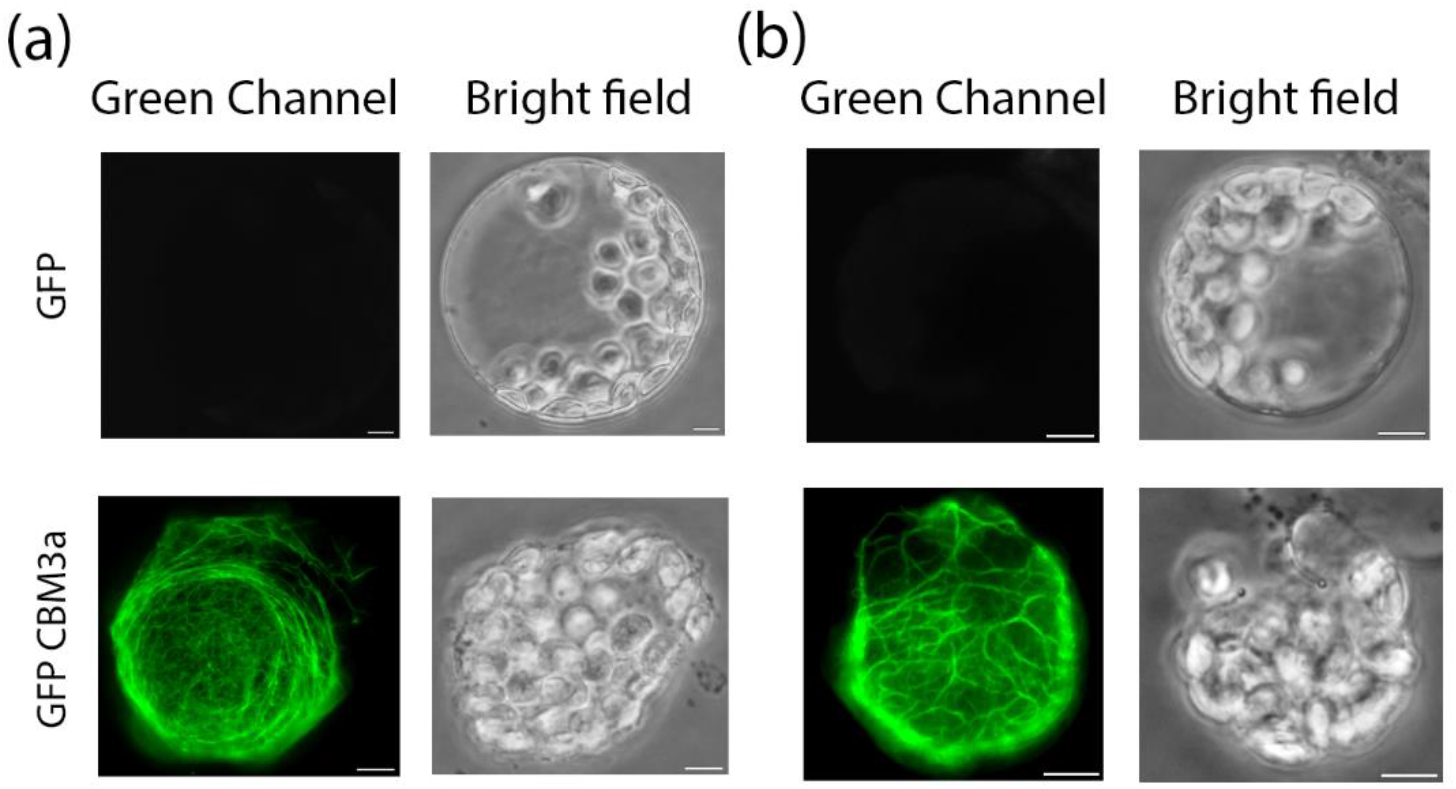
Fluorescence microscopy images of cell wall regenerated protoplasts. (a) GFP and GFP-CBM3a were added to the regenerated plant cell walls of Arabidopsis mesophyll protoplasts after 18 h incubation. (b) GFP and GFP-CBM3a were added along with the regeneration media at the outset of the 18 h incubation. Regenerated cellulose fibers labeled with GFP-CBM3a are visible under the green channel (left) and the bright field image of the intact protoplasts (right). (Scale bars, 6 μm.)

## Discussion

Carbohydrate binding modules (CBMs) are crucial in targeting plant cell wall polysaccharides in coordination with glycosyl hydrolases (Hervé *et al*., 2010; Fox *et al*., 2013). Several studies show the presence of tandem CBMs coexisting with these modular microbial plant cell wall glycosyl hydrolases (Freelove *et al*., 2001b; McCartney *et al*., 2006; Møller *et al*., 2021). Some of these tandem CBMs exhibit a synergistic effect with distinct specificities, and others exhibit an improved affinity towards polysaccharides. Unlike other plant polysaccharides that have an extensive repertoire of mAbs readily available, no stable mAb exists that can be used to visualize cellulose (Rydahl *et al*., 2018). Type-A CBMs bind specifically to cellulose, and particularly, CBM3a has a high affinity towards cellulose making it an appropriate probe to detect plant cellulose (Chowdhury *et al*., 2014; Johnsen *et al*., 2015). However, CBM3a could also binds non-specifically to other cell wall polysaccharides like xyloglucan (Hernandez-Gomez *et al*., 2015).

To address this issue and to increase the odds of CBM3a binding to cellulose with high affinity, we prepared tandem CBM3a fused to GFP to visualize the plant protoplasts directly. The ability of this tandem CBM3a to recognize isolated cellulose both *in vitro* and in regenerated plant cell walls was assessed. Interestingly, the tandem type-A CBMs – GFP-CBM3a and GFP-CBM64, showed reduced and equal affinity compared to their single domain counterparts. This reduction in binding affinity could be attributed to the native 42-aa linker that is highly flexible and might affect the binding of tandem CBMs. Furthermore, CBM3a exists as a dimer in nature and might dimerize because of this highly flexible linker leading to an increased desorption rate (Tormo *et al*., 1996; Arola & Linder, 2016). Interestingly, removing GFP seemed to decrease off-rate for tandem CBM3a drastically, suggesting that it might be more suitable for indirect immunofluorescence imaging purposes that involve a continuous and dynamic movement of the substrate in an aqueous environment. Additionally, these non-GFP-CBM versions could be conjugated to various amine-reactive fluorophores providing a library of CBMs across different wavelengths. The resulting fluorophore conjugated CBM proteins may exhibit greater photostability and brighter fluorescence than other fluorochromes (Moran-Mirabal *et al*., 2009).

In summary, these results provide qualitative and quantitative facets of the single and tandem CBMs used to imaging plant cellulose fibers. Although single CBMs show a reduced desorption rate from QCM-D assays, confocal imaging of plant protoplasts shows distinct cell walls with both single and tandem-CBMs. Linker length could play a critical role in binding irreversibly to the cellulose fibers, and usage of shorter and less flexible linkers might improve the binding affinity of tandem CBMs (Bandi *et al*., 2020). To unravel its complete potential, a more detailed study pertaining to the linker length of tandem CBMs for improving the binding affinity and specificity to cellulose in the regenerated plant cell wall is required. Furthermore, the plant cell wall is a natural amalgam of multiple polysaccharides intertwined with each other. Mimicking the exact composition of complex plant cell wall polysaccharides *in vitro* might be unfeasible. In order to overcome the non-specific binding nature exhibited by CBMs, techniques like phage display and directed evolution might be needed to engineer proteinaceous probes with greater specificity and affinity for imaging cellulose or other glycans synthesized and assembled in plant cell walls (DeVree *et al*., 2021). Lastly, an important characteristic of the GFP-CBM approach for detection and monitoring of cellulose fibers in plant cells is its apparent non-toxic nature. This property opens the possibility for direct visualization of the kinetic and spatial properties during the process of cell wall synthesis by using these protein-based labels for cellulose.

## Acknowledgements

The research was supported by the U.S. Department of Energy (Award Number: DE-SC0019313) and Rutgers University.

## Author Contributions

D.J. and S.P.S.C. designed the research. D.J., P.S., M.I., and J.S. conducted research. D.J. and P.S. analyzed the data. D.J. and S.P.S.C. wrote the manuscript with input from all authors.

## Data availability

All raw data used to support the findings of this study are available upon request from the corresponding author (Dr. Shishir P.S. Chundawat, Rutgers University).

## Supporting Information

**Fig. S1:** Plasmid maps of vectors and crystal structures of CBMs.

**Fig. S2:** Molecular architectures of CBM constructs.

**Fig. S3:** SDS-PAGE gel of purified CBMs and protein constructs.

**Fig. S4:** Langmuir-type adsorption model fits for different CBMs towards Cellulose-I.

**Fig. S5:** Langmuir-type adsorption model fits for different CBMs towards PASC.

**Fig. S6:** Binding Reversibility of CBMs towards Cellulose-I.

**Fig. S7:** Binding Reversibility of CBMs towards PASC.

**Fig. S8:** *K_off_* for non-GFP versions of CBM3a and CBM3a-CBM3a towards Cellulose-I.

**Fig. S9:** SDS-PAGE gel of purified CBMs and protein constructs.

**Fig. S10:** Fluorescence microscopy images of calcofluor white and GFP-CBM3a labeled plant protoplasts.

**Fig. S11:** Confocal laser scanning microscopy images of GFP-CBM64-CBM64 labeled plant protoplasts.

**Fig. S12:** Confocal laser scanning microscopy images of DSR-CBM labeled plant protoplasts.

**Supplementary video 1**: Confocal microscopy movie of regenerated Arabidopsis mesophyll protoplast in PBS buffer (no M2 media) with no GFP-CBM.

**Supplementary video 2**: Confocal microscopy movie of regenerated Arabidopsis mesophyll protoplast in M2 media with no GFP-CBM.

**Supplementary video 3**: Confocal microscopy movie of regenerated Arabidopsis mesophyll protoplast in M2 media labeled with GFP-CBM3a.

**Supplementary video 4**: Confocal microscopy movie of regenerated Arabidopsis mesophyll protoplast in M2 media labeled with GFP-CBM3a-CBM3a.

**Supplementary video 5**: Confocal microscopy movie of regenerated Arabidopsis mesophyll protoplast in M2 media labeled with GFP-CBM64-CBM64.

**Supplementary video 6**: Confocal microscopy movie of regenerated Arabidopsis mesophyll protoplast in M2 media labeled with DSR-CBM3a.

**Supplementary video 7**: Confocal microscopy movie of regenerated Arabidopsis mesophyll protoplast in M2 media labeled with DSR-CBM3a-CBM3a.

**Table S1:** Primer sequences used for cloning and sequencing of all constructs.

**Table S2:** Molecular weight and extinction coefficients of all CBMs used in this study.

**Supplementary Text S1:** Protein sequences of all major constructs used in this study.

