## Supplementary information for "Engineering and characterization of carbohydrate-binding modules to enable real-time imaging of cellulose fibrils biosynthesis in plant protoplasts"

**This SI file includes:**

Figures S1 to S11 (Pages 2-12)

Table S1 to S2 (Page 13)

Supplementary text S1 (Pages 14-15)

Supplementary video captions S1 to S7 (Page 16)

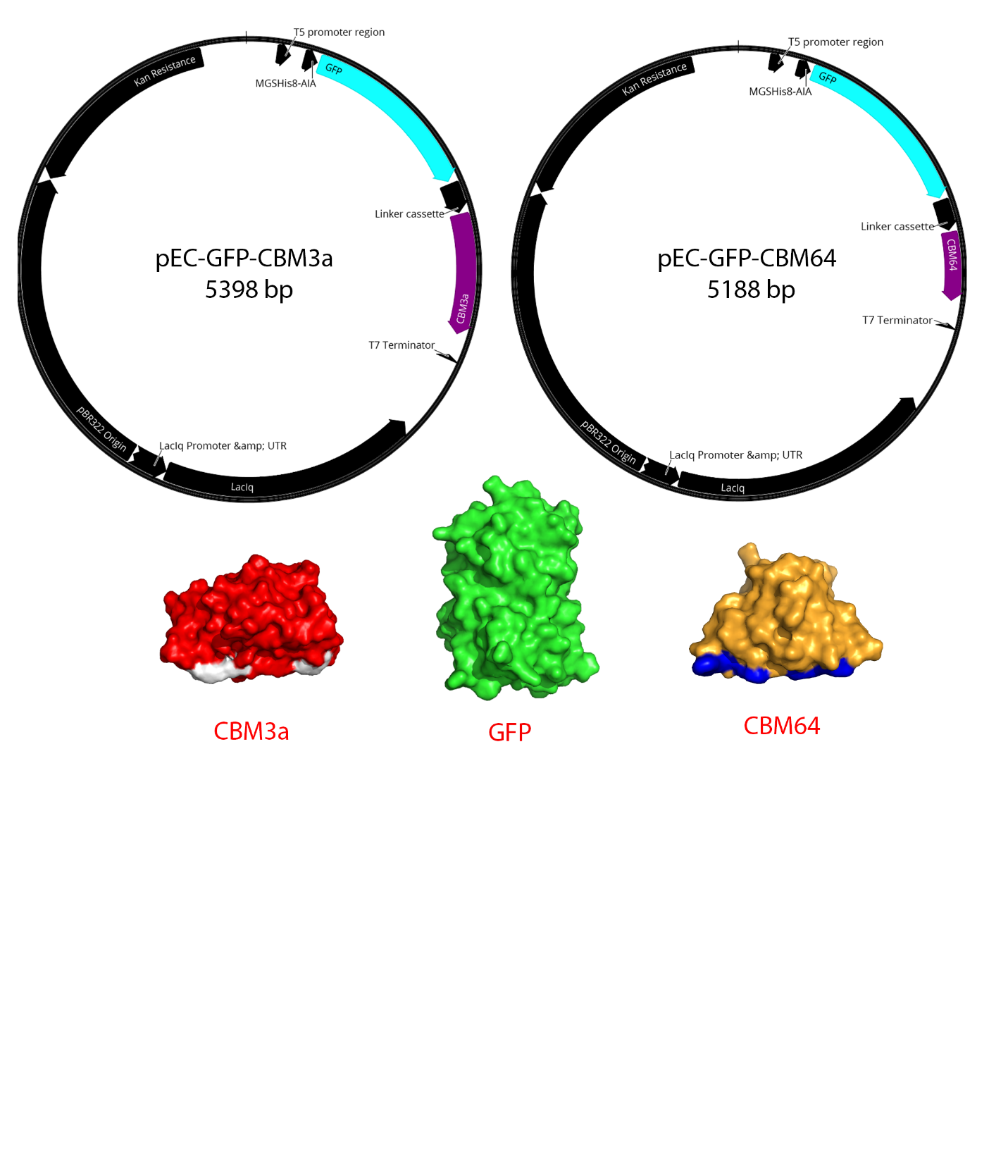

**Figure S1:** Plasmid maps for pEC-GFP-CBM3a and pEC-GFP-CBM64 indicating the T5 promoter region, 8x-HIS tag, followed by GFP-Linker-CBM3a and ending with T7 terminator. Crystal structures of CBM3a (*Clostridium thermocellum*, PDB ID: 1NBC), GFP(*Aequorea victoria*, PDB ID: 1GFL), and CBM64 (*Spirochaeta thermophila*, PDB ID: 5E9P) generated using PyMOL (bottom).

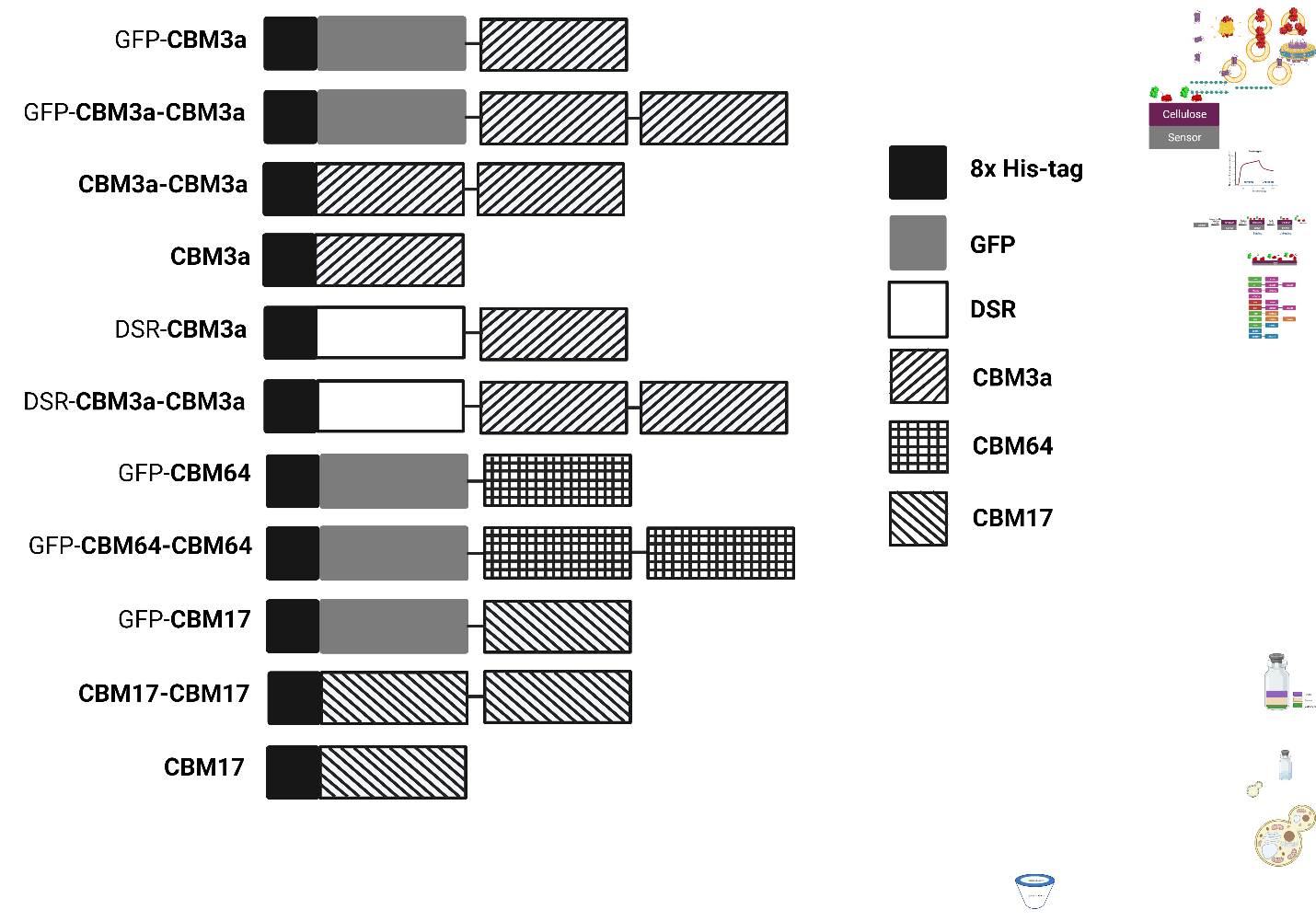

**Figure S2:** Molecular architectures of CBM constructs fused to GFP and other CBMs used in this study

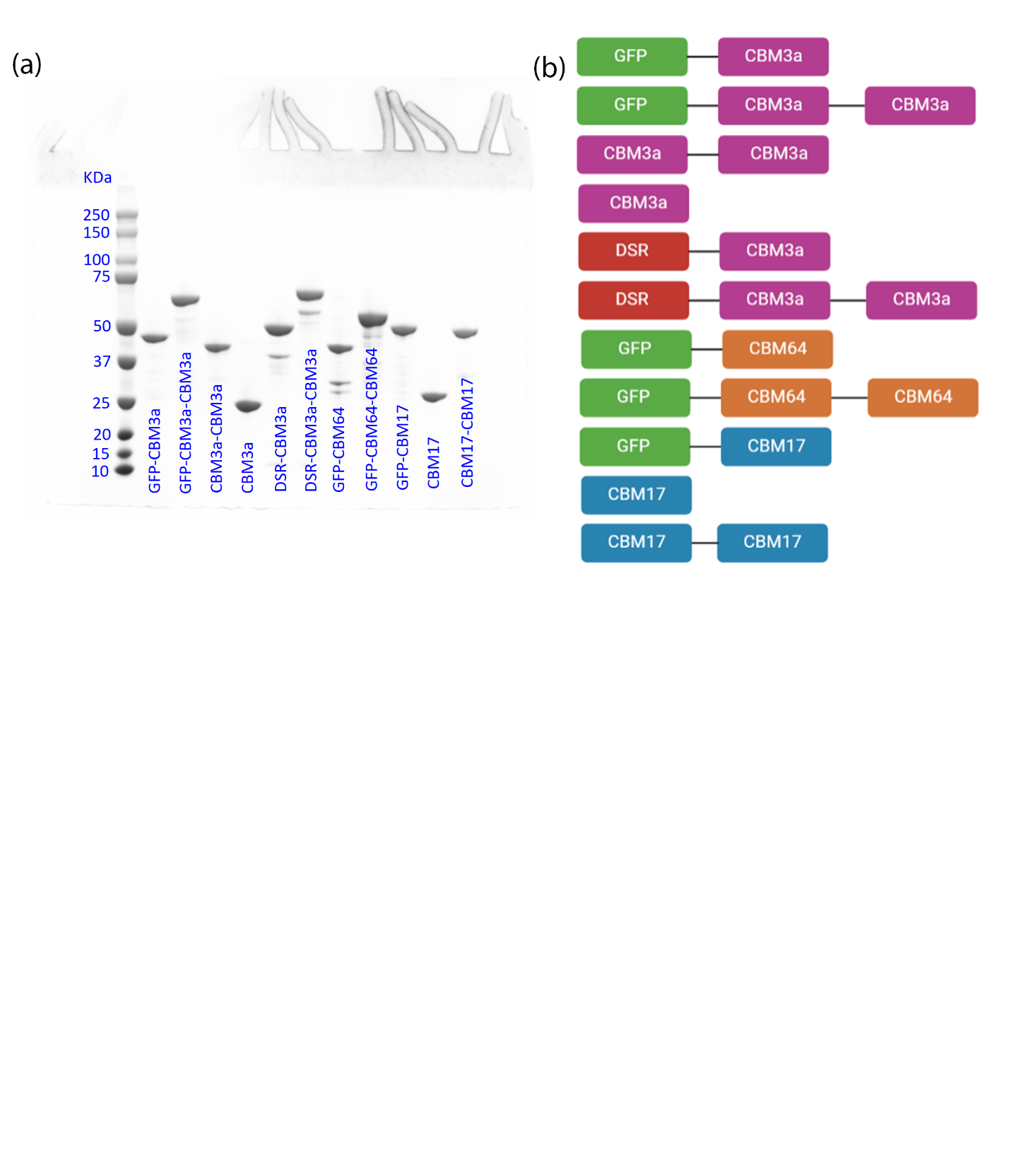

**Figure S3:** (a) SDS-PAGE gel for purified single and tandem CBM constructs; (b) different protein constructs used in this study

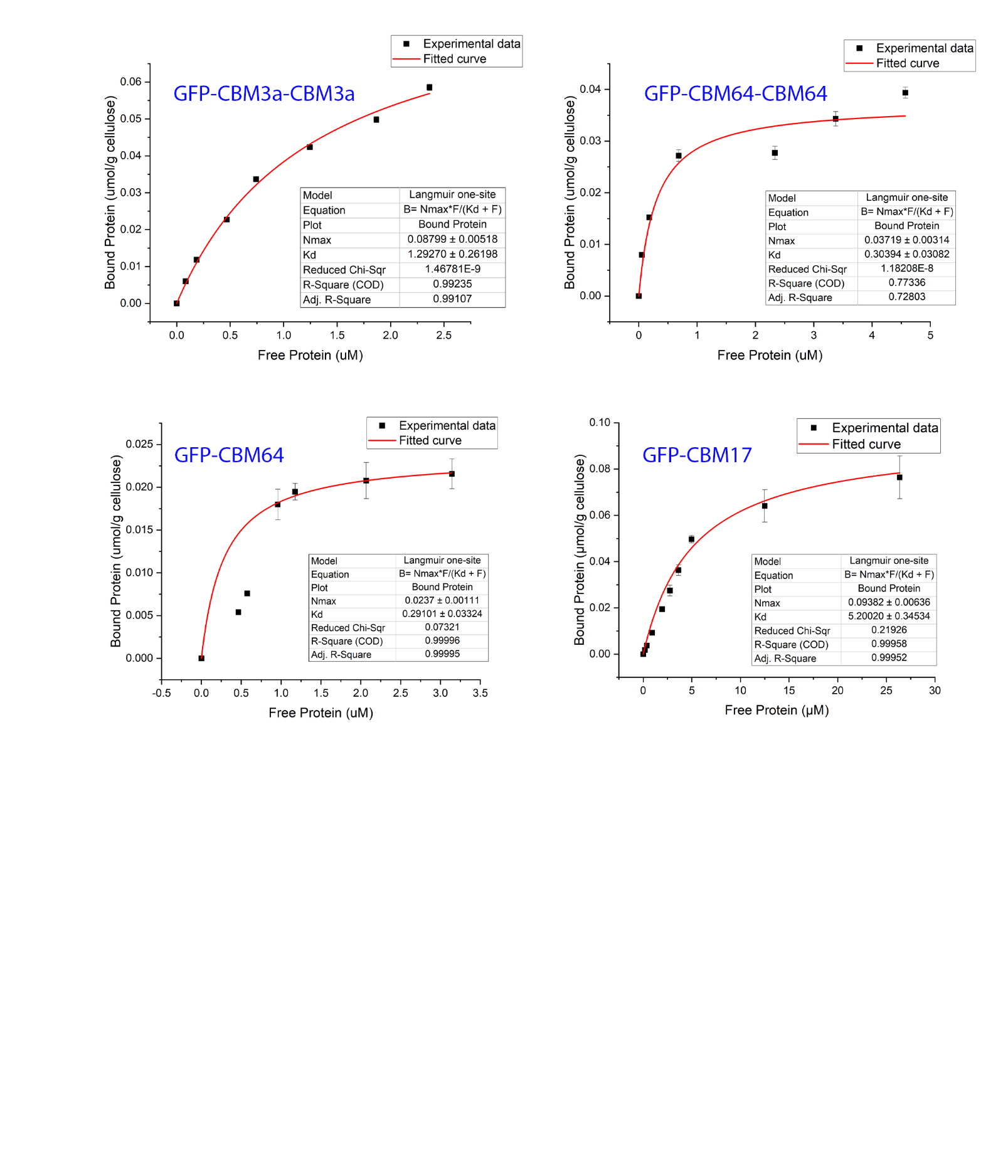

**Figure S4:** Langmuir-type adsorption model fits (in red) for different CBMs binding data (in black) to Avicel Cellulose I.

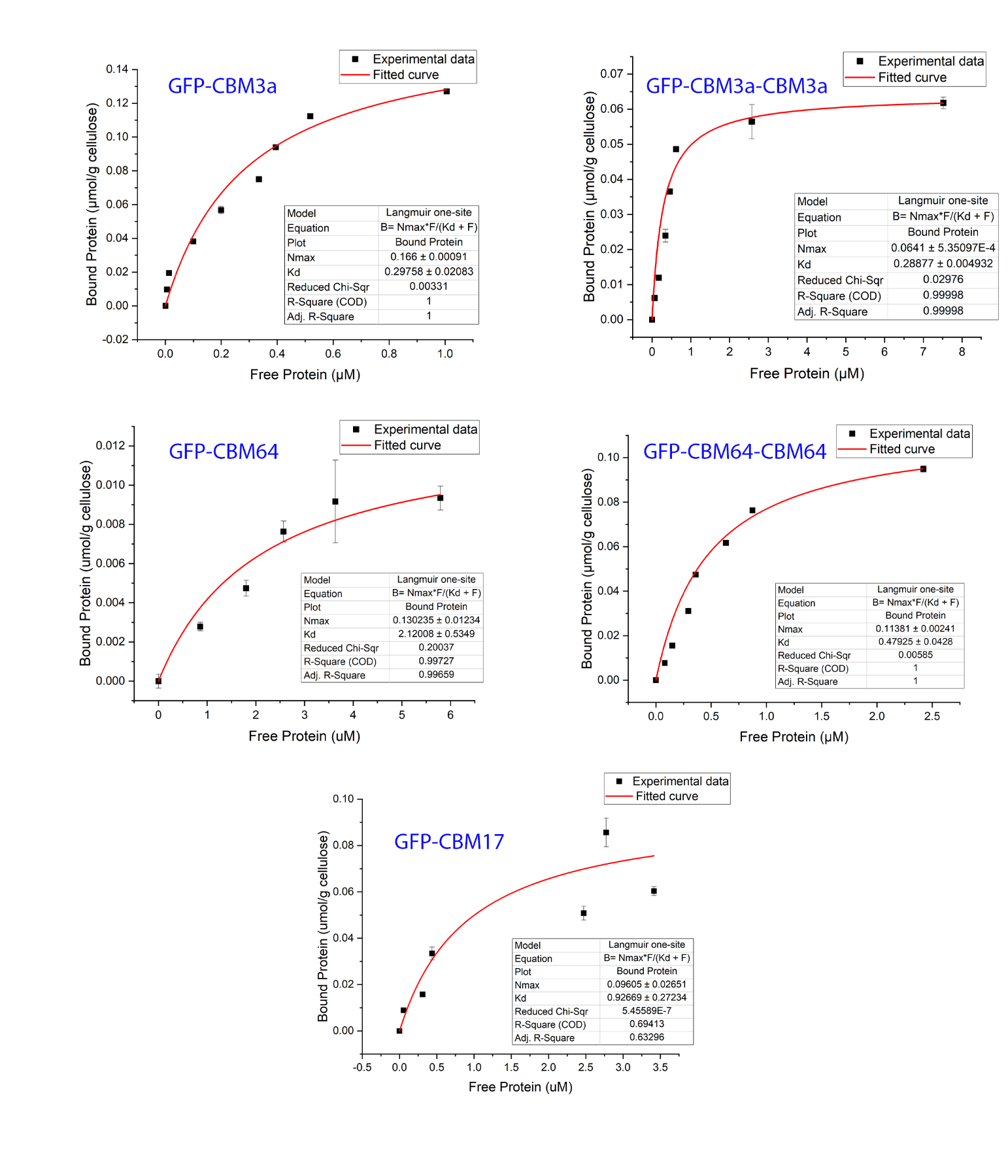

**Figure S5:** Langmuir-type adsorption model fits (in red) for different CBMs binding data (in black) to PASC.

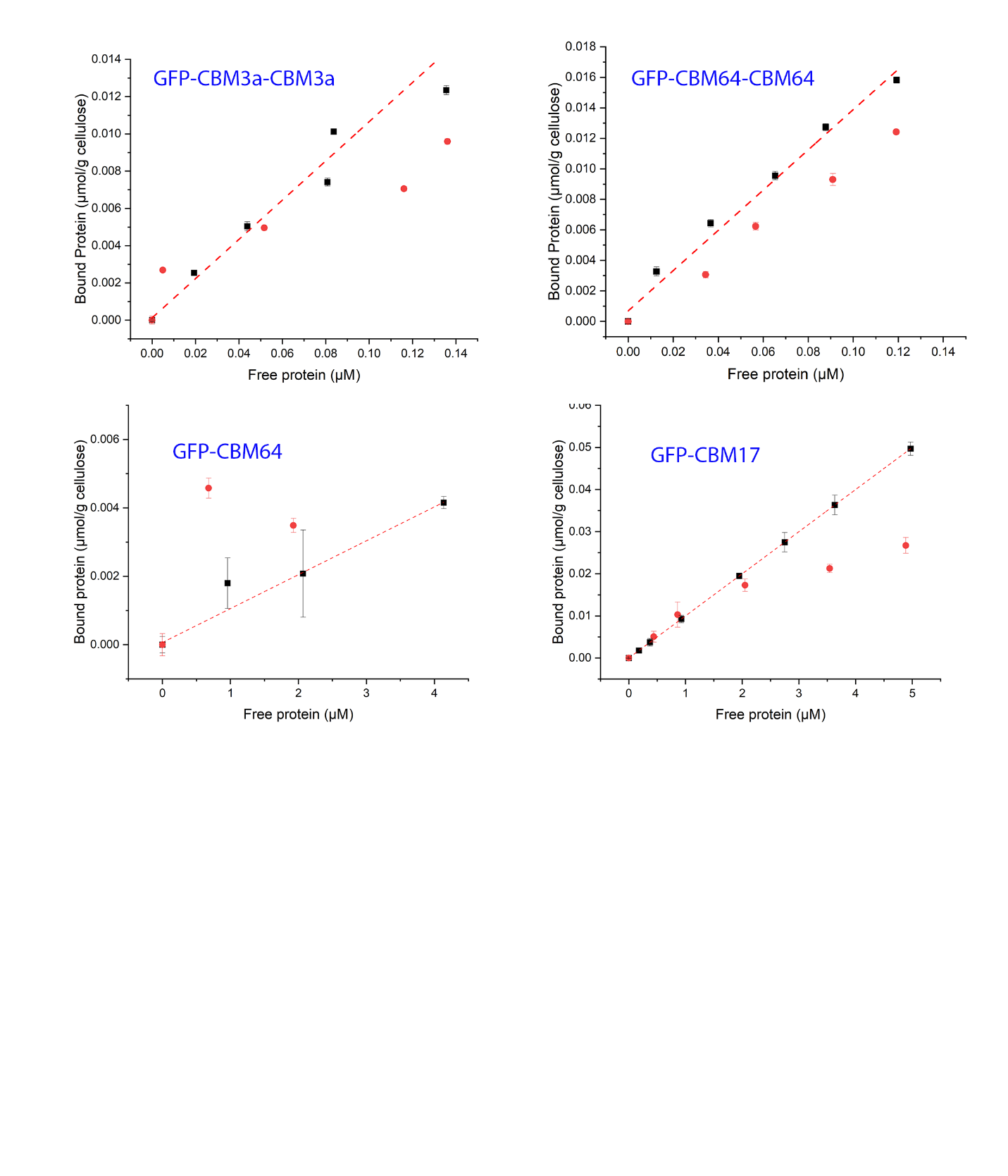

**Figure S6:** Raw data for binding assays to test reversibility of different CBMs binding to Avicel cellulose I. Error bars represent standard deviations based on at least three replicates and individual data points represent equilibrium reached at a given protein loading. Red dotted line represents linear fit joining original cellulose I data points respectively. Black squares and red circles represent data points obtained before and after re-equilibration for binding to cellulose I.

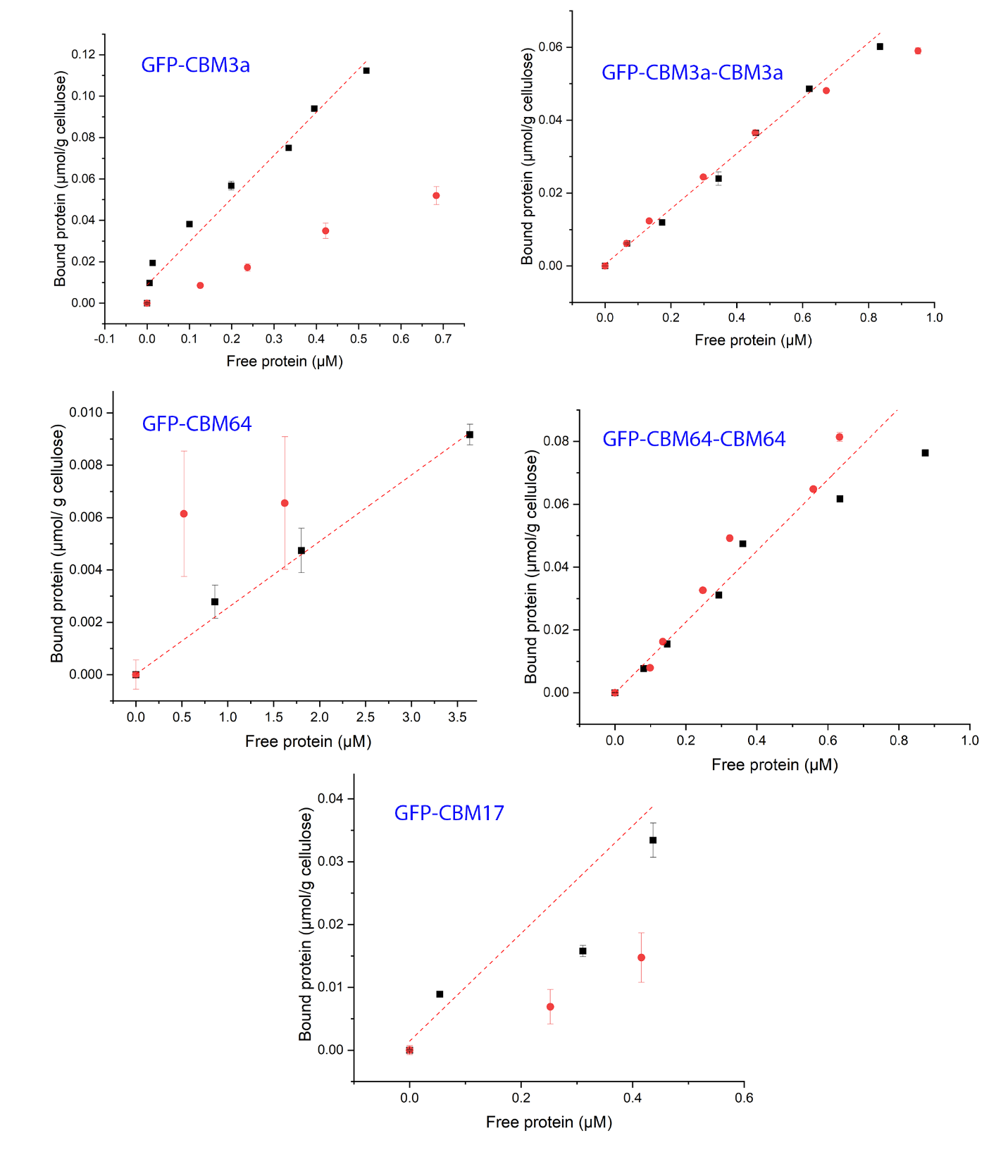

**Figure S7:** Raw data for binding assays to test reversibility of different CBMs binding to PASC. Error bars represent standard deviations based on at least three replicates and individual data points represent equilibrium reached at a given protein loading. Red dotted line represents linear fit joining original cellulose I data points respectively. Black squares and red circles represent data points obtained before and after re-equilibration for binding to cellulose I.

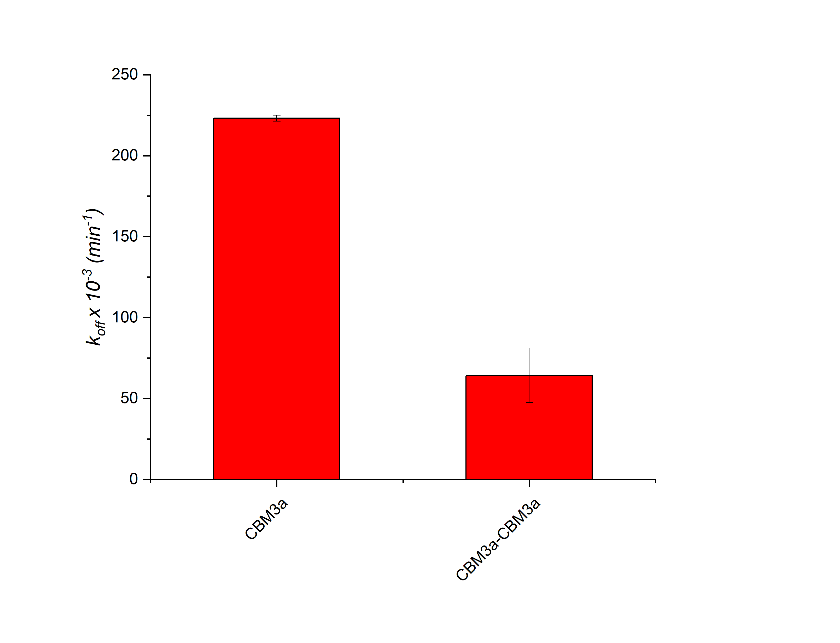

**Figure S8.** *K_off_* for non-GFP versions of CBM3a and CBM3a-CBM3a toward cellulose-I. CBM, carbohydrate‐binding module; GFP, green fluorescent protein.

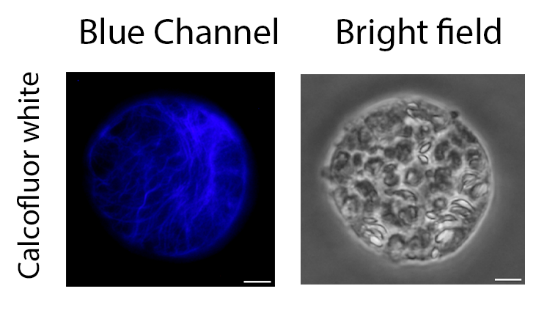

**A**

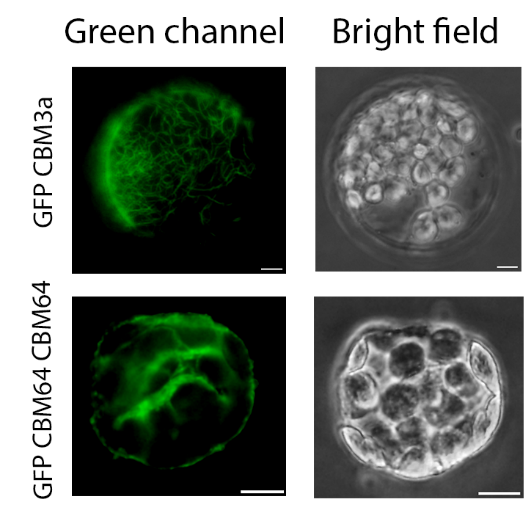

**B**

**Figure S9.** Fluorescence microscopy images of calcofluor white and GFP-CBM3a bound to cellulose in regenerated plant cell walls of Arabidopsis mesophyll protoplasts. Regenerated cellulose fibers are visible under the blue channel for calcofluor white (left) and bright field view (right) (A) ; cellulose fibers are visible under the green channel for GFP-CBM3a (left) and bright field view (right) (B) (Scale bars, 6 µm.)

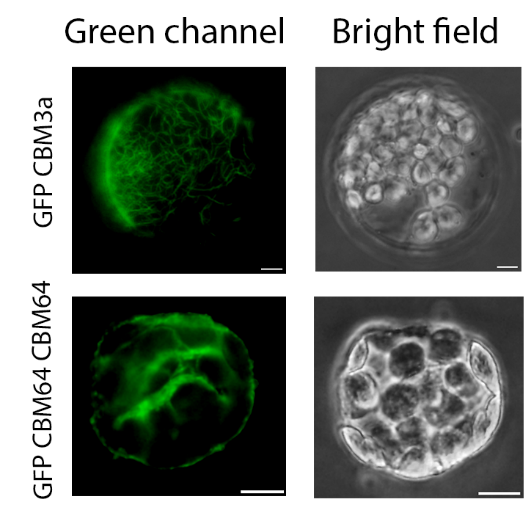

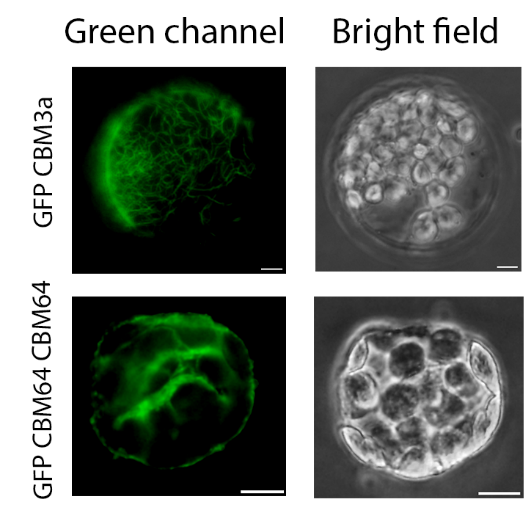

**Figure S10.** Confocal laser scanning microscopy images of GFP-CBM64-CBM64 bound to cellulose in regenerated plant cell walls of Arabidopsis mesophyll protoplasts. Regenerated cellulose fibers are visible under the green channel (left) and bright field view is shown on the right (Scale bars, 6 µm.)

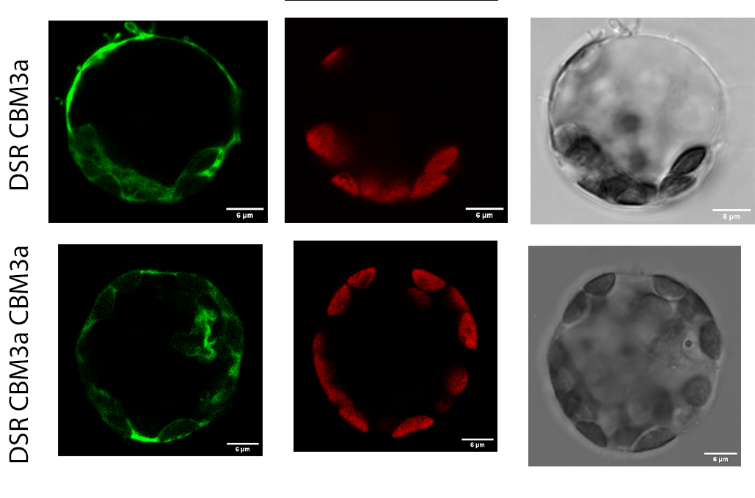

**Figure S11.** Confocal laser scanning microscopy images of DSR-CBM3a and DSR-CBM3a-CBM3a bound to regenerated plant cell walls of Arabidopsis mesophyll protoplasts (left), autofluorescence from chlorophyll of the plastids (middle), and bright field view (right). The color of the DSR labeled cell wall has been changed to green manually using ImageJ. (Scale bars, 6 µm.)

**Table S1**. Forward (FP) and reverse (RP) primer sequences for performing SLIC to generate all constructs reported in this paper. The last two primer sequences were used for confirming sequences.

| **Name of Primer** | **Sequence** |
| --- | --- |
| Vec_For_GFP_CBM3a_CBM3a | AGAACCAGGATAAGGATCCTCTAGAGTC |
| Vec_Rev_GFP_CBM3a_CBM3a | GCGTTTAAACCTCCTGGTTCTTTCCCCC |
| Ins_For_Linker_CBM3a_CBM3a | AGAACCAGGAGGTTTAAACGCGACTCCC |
| Ins_Rev_Linker_CBM3a_CBM3a | AGGATCCTTATCCTGGTTCTTTCCCCCA |
| Vec_For_GFP_CBM64_CBM64 | GTGGAAATCAAATAAGGATCCTCTAGAGTCG |
| Vec_Rev_GFP_CBM64_CBM64 | CGTTTAAACCTTTGATTTCCACATGCGA |
| Ins_For_Linker_CBM64_CBM64 | GGAAATCAAAGGTTTAAACGCGACTCCC |
| Ins_Rev_Linker_CBM64_CBM64 | AGGATCCTTATTTGATTTCCACATGCGA |
| Vec_For_CBM3a_CBM3a | AAGAACCAGGATAAGGATCCTCTAGAGTC |
| Vec_Rev_CBM3a_CBM3a | CGTTTAAACCCTGGAAGTACAGGTTTTC |
| Ins_For_CBM3a_CBM3a | GTACTTCCAGGGTTTAAACGCGACTCCC |
| Ins_Rev_CBM3a_CBM3a | AGGATCCTTATCCTGGTTCTTTCCCCCA |
| For_ CBM3a | AGAACCAGGATAAGGATCCTCTAGAGTC |
| Rev_CBM3a | AGGATCCTTATCCTGGTTCTTTCCCCCA |
| Vec_For_DSR_CBM3a_CBM3a | AGAACCAGGATAAGGATCCTCTAGAGTC |
| Vec_Rev_DSR_CBM3a_CBM3a | CGTTTAAACCTCCTGGTTCTTTCCCCCA |
| Ins_For_ DSR_CBM3a_CBM3a | AGAACCAGGAGGTTTAAACGCGACTCCC |
| Ins_Rev_ DSR_CBM3a_CBM3a | AGGATCCTTATCCTGGTTCTTTCCCCCA |
| Vec_For_CBM17_CBM17 | ATTTACCAAAGGTTTAAACGCGACTCCC |
| Vec_Rev_CBM17_CBM17 | TGGGAGTCGCGGCGATCGCCTGGAAGTA |
| Ins_For_CBM17_CBM17 | GGCGATCGCCGCGACTCCCACTAAAGGT |
| Ins_Rev_CBM17_CBM17 | CGTTTAAACCTTTGGTAAATTTGATGTT |
| CBM17_For | GGCGATCGCCGGTTTAAACGCGACTCCC |
| CBM17_Rev | CGTTTAAACCGGCGATCGCCTGGAAGTA |
| Sequencing_Forward | AAATAGGGGTTCCGCG |
| Sequencing_Reverse | AGAAACGAGTTTGCTAGT |

**Table S2**. Molecular weight and extinction coefficients of all proteins reported in this paper

| **Name of Protein** | **Molecular weight (Da)** | **Extinction coefficient (M^-1^ cm^-1^)** |
| --- | --- | --- |
| GFP-CBM3a | 51053.65 | 58915 |
| GFP-CBM3a-CBM3a | 72996.69 | 94450 |
| CBM3a-CBM3a | 46142.49 | 72435 |
| CBM3a | 24199.45 | 36900 |
| DSR-CBM3a | 48966.17 | 71405 |
| DSR-CBM3a-CBM3a | 71809.21 | 106815 |
| GFP-CBM64 | 43941.81 | 71405 |
| GFP-CBM64-CBM64 | 58773.01 | 119305 |
| GFP-CBM17 | 52930.82 | 53985 |
| CBM17 | 26331.94 | 31970 |
| CBM17-CBM17 | 49867.84 | 62450 |

**Supplementary Text S1.** Protein sequences of all major constructs used in this study.

>>GFP-CBM3a-WT

MGHHHHHHHHASENLYFQAIASKGEELFTGVVPILVELDGDVNGHKFSVSGEGEGDATYGKLTLKFICTTGKLPVPWPTLVTTLTYGVQCFSRYPDHMKQHDFFKSAMPEGYVQERTISFKDDGNYKTRAEVKFEGDTLVNRIELKGIDFKEDGNILGHKLEYNYNSHNVYITADKQKNGIKANFKIRHNIEDGSVQLADHYQQNTPIGDGPVLLPDNHYLSTQSALSKDPNEKRDHMVLLEFVTAAGITHGMDELYKGLNATPTKGATPTNTATPTKSATATPTRPSVPTNTPTNTPANTLKVSGNLKVEFYNSNPSDTTNSINPQFKVTNTGSSAIDLSKLTLRYYYTVDGQKDQTFWCDHAAIIGSNGSYNGITSNVKGTFVKMSSSTNNADTYLEISFTGGTLEPGAHVQIQGRFAKNDWSNYTQSNDYSFKSASQFVEWDQVTAYLNGVLVWGKEPG

>>GFP-CBM3a-CBM3a

MGHHHHHHHHASENLYFQAIASKGEELFTGVVPILVELDGDVNGHKFSVSGEGEGDATYGKLTLKFICTTGKLPVPWPTLVTTLTYGVQCFSRYPDHMKQHDFFKSAMPEGYVQERTISFKDDGNYKTRAEVKFEGDTLVNRIELKGIDFKEDGNILGHKLEYNYNSHNVYITADKQKNGIKANFKIRHNIEDGSVQLADHYQQNTPIGDGPVLLPDNHYLSTQSALSKDPNEKRDHMVLLEFVTAAGITHGMDELYKGLNATPTKGATPTNTATPTKSATATPTRPSVPTNTPTNTPANTLKVSGNLKVEFYNSNPSDTTNSINPQFKVTNTGSSAIDLSKLTLRYYYTVDGQKDQTFWCDHAAIIGSNGSYNGITSNVKGTFVKMSSSTNNADTYLEISFTGGTLEPGAHVQIQGRFAKNDWSNYTQSNDYSFKSASQFVEWDQVTAYLNGVLVWGKEPGGLNATPTKGATPTNTATPTKSATATPTRPSVPTNTPTNTPANTLKVSGNLKVEFYNSNPSDTTNSINPQFKVTNTGSSAIDLSKLTLRYYYTVDGQKDQTFWCDHAAIIGSNGSYNGITSNVKGTFVKMSSSTNNADTYLEISFTGGTLEPGAHVQIQGRFAKNDWSNYTQSNDYSFKSASQFVEWDQVTAYLNGVLVWGKEPG

>>DSR-CBM3a

MGHHHHHHHHASENLYFQAIAMDNTEDVIKEFMQFKVRMEGSVNGHYFEIEGEGEGKPYEGTQTAKLQVTKGGPLPFAWDILSPQFQYGSKAYVKHPADIPDYMKLSFPEGFTWERSMNFEDGGVVEVQQDSSLQDGTFIYKVKFKGVNFPADGPVMQKKTAGWEPSTEKLYPQDGVLKGEISHALKLKDGGHYTCDFKTVYKAKKPVQLPGNHYVDSKLDITNHNEDYTVVEQYEHAEARHSGSQGLNATPTKGATPTNTATPTKSATATPTRPSVPTNTPTNTPANTLKVSGNLKVEFYNSNPSDTTNSINPQFKVTNTGSSAIDLSKLTLRYYYTVDGQKDQTFWCDHAAIIGSNGSYNGITSNVKGTFVKMSSSTNNADTYLEISFTGGTLEPGAHVQIQGRFAKNDWSNYTQSNDYSFKSASQFVEWDQVTAYLNGVLVWGKEPG

>>DSR-CBM3a-CBM3a

MGHHHHHHHHASENLYFQAIAMDNTEDVIKEFMQFKVRMEGSVNGHYFEIEGEGEGKPYEGTQTAKLQVTKGGPLPFAWDILSPQFQYGSKAYVKHPADIPDYMKLSFPEGFTWERSMNFEDGGVVEVQQDSSLQDGTFIYKVKFKGVNFPADGPVMQKKTAGWEPSTEKLYPQDGVLKGEISHALKLKDGGHYTCDFKTVYKAKKPVQLPGNHYVDSKLDITNHNEDYTVVEQYEHAEARHSGSQGLNATPTKGATPTNTATPTKSATATPTRPSVPTNTPTNTPANTLKVSGNLKVEFYNSNPSDTTNSINPQFKVTNTGSSAIDLSKLTLRYYYTVDGQKDQTFWCDHAAIIGSNGSYNGITSNVKGTFVKMSSSTNNADTYLEISFTGGTLEPGAHVQIQGRFAKNDWSNYTQSNDYSFKSASQFVEWDQVTAYLNGVLVWGKEPGGLNATPTKGATPTNTATPTKSATATPTRPSVPTNTPTNTPANTLKVSGNLKVEFYNSNPSDTTNSINPQFKVTNTGSSAIDLSKLTLRYYYTVDGQKDQTFWCDHAAIIGSNGSYNGITSNVKGTFVKMSSSTNNADTYLEISFTGGTLEPGAHVQIQGRFAKNDWSNYTQSNDYSFKSASQFVEWDQVTAYLNGVLVWGKEPG

>>CBM3a

MGHHHHHHHHASENLYFQAIAVSGNLKVEFYNSNPSDTTNSINPQFKVTNTGSSAIDLSKLTLRYYYTVDGQKDQTFWCDHAAIIGSNGSYNGITSNVKGTFVKMSSSTNNADTYLEISFTGGTLEPGAHVQIQGRFAKNDWSNYTQSNDYSFKSASQFVEWDQVTAYLNGVLVWGKEPG

>>CBM3a-CBM3a

MGHHHHHHHHASENLYFQGLNATPTKGATPTNTATPTKSATATPTRPSVPTNTPTNTPANTLKVSGNLKVEFYNSNPSDTTNSINPQFKVTNTGSSAIDLSKLTLRYYYTVDGQKDQTFWCDHAAIIGSNGSYNGITSNVKGTFVKMSSSTNNADTYLEISFTGGTLEPGAHVQIQGRFAKNDWSNYTQSNDYSFKSASQFVEWDQVTAYLNGVLVWGKEPGGLNATPTKGATPTNTATPTKSATATPTRPSVPTNTPTNTPANTLKVSGNLKVEFYNSNPSDTTNSINPQFKVTNTGSSAIDLSKLTLRYYYTVDGQKDQTFWCDHAAIIGSNGSYNGITSNVKGTFVKMSSSTNNADTYLEISFTGGTLEPGAHVQIQGRFAKNDWSNYTQSNDYSFKSASQFVEWDQVTAYLNGVLVWGKEPG

>>GFP-CBM64-WT

MGHHHHHHHHASENLYFQAIASKGEELFTGVVPILVELDGDVNGHKFSVSGEGEGDATYGKLTLKFICTTGKLPVPWPTLVTTLTYGVQCFSRYPDHMKQHDFFKSAMPEGYVQERTISFKDDGNYKTRAEVKFEGDTLVNRIELKGIDFKEDGNILGHKLEYNYNSHNVYITADKQKNGIKANFKIRHNIEDGSVQLADHYQQNTPIGDGPVLLPDNHYLSTQSALSKDPNEKRDHMVLLEFVTAAGITHGMDELYKGLNATPTKGATPTNTATPTKSATATPTRPSVPTNTPTNTPANTLKPTPSGEYTAIALPFTYDGAGEYYWKTDQFSTDPNDWSRYVNSWNLDLLEINGTDYTNVWVAQHQITPASDGYWYIHYKGSYPWSHVEIK

>>GFP-CBM64-CBM64

MGHHHHHHHHASENLYFQAIASKGEELFTGVVPILVELDGDVNGHKFSVSGEGEGDATYGKLTLKFICTTGKLPVPWPTLVTTLTYGVQCFSRYPDHMKQHDFFKSAMPEGYVQERTISFKDDGNYKTRAEVKFEGDTLVNRIELKGIDFKEDGNILGHKLEYNYNSHNVYITADKQKNGIKANFKIRHNIEDGSVQLADHYQQNTPIGDGPVLLPDNHYLSTQSALSKDPNEKRDHMVLLEFVTAAGITHGMDELYKGLNATPTKGATPTNTATPTKSATATPTRPSVPTNTPTNTPANTLKPTPSGEYTAIALPFTYDGAGEYYWKTDQFSTDPNDWSRYVNSWNLDLLEINGTDYTNVWVAQHQITPASDGYWYIHYKGSYPWSHVEIKGLNATPTKGATPTNTATPTKSATATPTRPSVPTNTPTNTPANTLKPTPSGEYTAIALPFTYDGAGEYYWKTDQFSTDPNDWSRYVNSWNLDLLEINGTDYTNVWVAQHQITPASDGYWYIHYKGSYPWSHVEIK

>>GFP-CBM17-WT

MGHHHHHHHHASENLYFQAIASKGEELFTGVVPILVELDGDVNGHKFSVSGEGEGDATYGKLTLKFICTTGKLPVPWPTLVTTLTYGVQCFSRYPDHMKQHDFFKSAMPEGYVQERTISFKDDGNYKTRAEVKFEGDTLVNRIELKGIDFKEDGNILGHKLEYNYNSHNVYITADKQKNGIKANFKIRHNIEDGSVQLADHYQQNTPIGDGPVLLPDNHYLSTQSALSKDPNEKRDHMVLLEFVTAAGITHGMDELYKGLNATPTKGATPTNTATPTKSATATPTRPSVPTNTPTNTPANTLKSQPTAPKDFSSGFWDFNDGTTQGFGVNPDSPITAINVENANNALKISNLNSKGSNDLSEGNFWANVRISADIWGQSINIYGDTKLTMDVIAPTPVNVSIAAIPQSSTHGWGNPTRAIRVWTNNFVAQTDGTYKATLTISTNDSPNFNTIATDAADSVVTNMILFVGSNSDNISLDNIKFTK

**Supplementary video 1**: Confocal microscopy movie of regenerated Arabidopsis mesophyll protoplast in PBS buffer (no M2 media) with no GFP-CBM.

**Supplementary video 2**: Confocal microscopy movie of regenerated Arabidopsis mesophyll protoplast in M2 media with no GFP-CBM.

**Supplementary video 3**: Confocal microscopy movie of regenerated Arabidopsis mesophyll protoplast in M2 media labeled with GFP-CBM3a.

**Supplementary video 4**: Confocal microscopy movie of regenerated Arabidopsis mesophyll protoplast in M2 media labeled with GFP-CBM3a-CBM3a.

**Supplementary video 5**: Confocal microscopy movie of regenerated Arabidopsis mesophyll protoplast in M2 media labeled with GFP-CBM64-CBM64.

**Supplementary video 6**: Confocal microscopy movie of regenerated Arabidopsis mesophyll protoplast in M2 media labeled with DSR-CBM3a.

**Supplementary video 7**: Confocal microscopy movie of regenerated Arabidopsis mesophyll protoplast in M2 media labeled with DSR-CBM3a-CBM3a.
